# Behavioral and Neural Variability of Naturalistic Arm Movements

**DOI:** 10.1101/2020.04.17.047357

**Authors:** Steven M. Peterson, Satpreet H. Singh, Nancy X. R. Wang, Rajesh P. N. Rao, Bingni W. Brunton

## Abstract

Motor behaviors are central to many functions and dysfunctions of the brain, and understanding their neural basis has consequently been a major focus in neuroscience. However, most studies of motor behaviors have been restricted to artificial, repetitive paradigms, far removed from natural movements performed “in the wild.” Here, we leveraged recent advances in machine learning and computer vision to analyze intracranial recordings from 12 human subjects during thousands of spontaneous, unstructured arm reach movements, observed over several days for each subject. These naturalistic movements elicited cortical spectral power patterns consistent with findings from controlled paradigms, but with considerable neural variability across subjects and events. We modeled inter-event variability using ten behavioral and environmental features; the most important features explaining this variability were reach angle and day of recording. Our work is among the first studies connecting behavioral and neural variability across cortex in humans during unstructured movements and contributes to our understanding of long-term naturalistic behavior.

---

Natural human movements are remarkable in their complexity and adaptability, relying on precisely coordinated sensorimotor processing in several cortical regions [1–4]. Much of our understanding on the neural basis of human upper-limb movements has been gained by studying constrained, repetitive movements in the laboratory, using paradigms such as the center-out reaching task [5–9]. Center-out reaching is an elegant method for investigating the neural basis of movement, but it remains unclear how well its findings generalize to the spontaneous, unstructured actions observed in the real world [10, 11]. Studies have enhanced the realism of experimental reaching paradigms by incorporating self-cued and less restrictive movements [12–15], but few studies have focused on completely unstructured, naturalistic human movements recorded outside of defined laboratory paradigms. Focusing on such naturalistic behavior enriches our understanding of the relationship between motor behavior and cortical activation [16, 17] and motivates development of robust brain-computer interfaces to restore impaired movement and sensation across diverse contexts [18–22].

The history of modern neuroscience has seen a consistent trend towards studies incorporating more naturalistic elements. The use of stimuli, environments, and tasks with increasing ecological relevance to the animal has enhanced our understanding of how the brain functions, complementing results from more artificial laboratory paradigms. For instance, early neural recordings focused on anesthetized animals, but the transition to experiments with awake, behaving animals transformed our knowledge of sensory, motor, and cognitive brain functions [23–27]. More recently, researchers have moved towards using natural auditory and visual stimuli, finding novel neural responses not seen with artificial stimuli [28–33]; moreover, features of natural stimuli often better explain the observed variance in neural activity [34, 35]. In human neuroscience and behavior, advances in technology have enabled an expanded focus on a variety of mobile outdoor paradigms [36–40], spatial navigation tasks within immersive virtual environments [41, 42], and tasks involving active social interactions [43–46].

Intracranial electrophysiological recordings offer a unique view into the neural correlates of human behavior. These recordings, obtained using electrocorticography (ECoG), contain physiologically relevant spectral power patterns corresponding to a variety of behaviors [47–51]. ECoG recording electrodes are implanted on the cortical surface, beneath the skull and dura; these signals are thus cleaner and less susceptible to artifact contamination than signals from electroencephalography (EEG) [52]. Although implanting ECoG electrodes is an invasive neurosurgical procedure, the recordings are highly informative and have a combination of high spatial and temporal resolution not found in other human neuroimaging or neural recording modalities [53–55].

During instructed upper limb movements, ECoG spectral power in fronto-parietal cortical areas, particularly over sensorimotor cortex, has been shown to transiently increase at high frequencies and decrease at low frequencies [4, 56–58]. Similar spectral power changes have also been observed in EEG and local field potential recordings across a wide variety of movement behaviors [59–63]. An important attribute of ECoG recordings is that the patients are being continuously monitored over long periods of time, often approximately a week, providing unique opportunities to collect long-term datasets during unconstrained, uninstructed movements [64–69]. However, the behavioral and neural variability of such spontaneous, naturalistic movements remains unexplored.

Analyzing naturalistic data presents formidable challenges, but recent innovations in data science make it possible to extract meaningful findings from increasingly complex, including naturalistic and opportunistic, datasets [72]. Without prior experimental design or direct behavioral measurements, a critical first step in analyzing naturalistic data had previously been laborious manual annotation of behavior. Such tedious labeling severely limits the amount of usable data and is prone to subjective error. Fortunately, recent advances in computer vision and machine learning have enabled substantial automation of the analysis and quantification of naturalistic behaviors [70, 73–76]. Even so, making sense of annotated behavior remains challenging in the absence of a controlled experimental paradigm. There are often many possible ways to characterize behavioral features, making it challenging to select ones that are objective and neurally-relevant. Fortunately, previous upper-limb movement studies have identified several neurally-relevant behavioral features that can be obtained without subjective, manual identification. Such behavioral features include the angle, duration, magnitude, and velocity of the movement as well as whether or not the movement was bimanual [77–80]. Based on previous research [81, 82], we were also motivated to consider the effects of social interactions on neural activity. Finally, ECoG recordings are non-stationary [83–85], so we considered on how movement-related neural activity varied over several hours and across recording days.

In this paper, we analyzed opportunistic, clinical intracranial recordings from 12 human subjects across 3–5 days each as we observed their naturalistic spontaneous arm movements. We developed an automated approach to identify and characterize thousands of spontaneous arm movements, enabling scal-able analysis of video that was acquired simultaneously with the intracranial recordings. We characterized the variability of both naturalistic upper-limb reaching movements and the corresponding changes in cortical spectral power. Based on findings from controlled experiments, we hypothesized that naturalistic reaches would be associated with transient decreases in low-frequency power and increases in high-frequency power, localized to fronto-parietal sensorimotor cortices [4, 56]. Our results support this hypothesis on average; however, we show that there is considerable variability in spectral power both within and across subjects. Multiple-variable linear regression modelling partially explains this single-event neural variability using reach angle and day of recording features, but much of the neural variability remains unexplained by behavioral and environmental features. To support reproducibility and facilitate future research in naturalistic human movement analysis, we have publicly released our curated dataset containing synchronized behavioral and neural data.

## Results

We describe behavioral and neural variability observed in multielectrode intracranial neural recordings and video from 12 human subjects during thousands of unstructured arm movements. Each subject had been implanted with electrocorticography (ECoG) electrodes for clinical monitoring, and we analyzed 3–5 days of simultaneously recorded video and electrophysiological data following surgery. We developed an automated and scalable approach to track upper limb movements based on machine learning and then focused on analyzing spectral power changes associated with movements of the wrist contralateral to the hemisphere with implanted electrodes (Fig. 1a–g). ECoG monitoring was clinically motivated, so there was substantial variation in electrode placement among subjects (Fig. 1h). Because our focus was on arm reaching behavior, we chose to analyze 12 subjects who were generally active during their monitoring and also had electrodes implanted over fronto-parietal sensorimotor cortical areas.

**Fig. 1:**
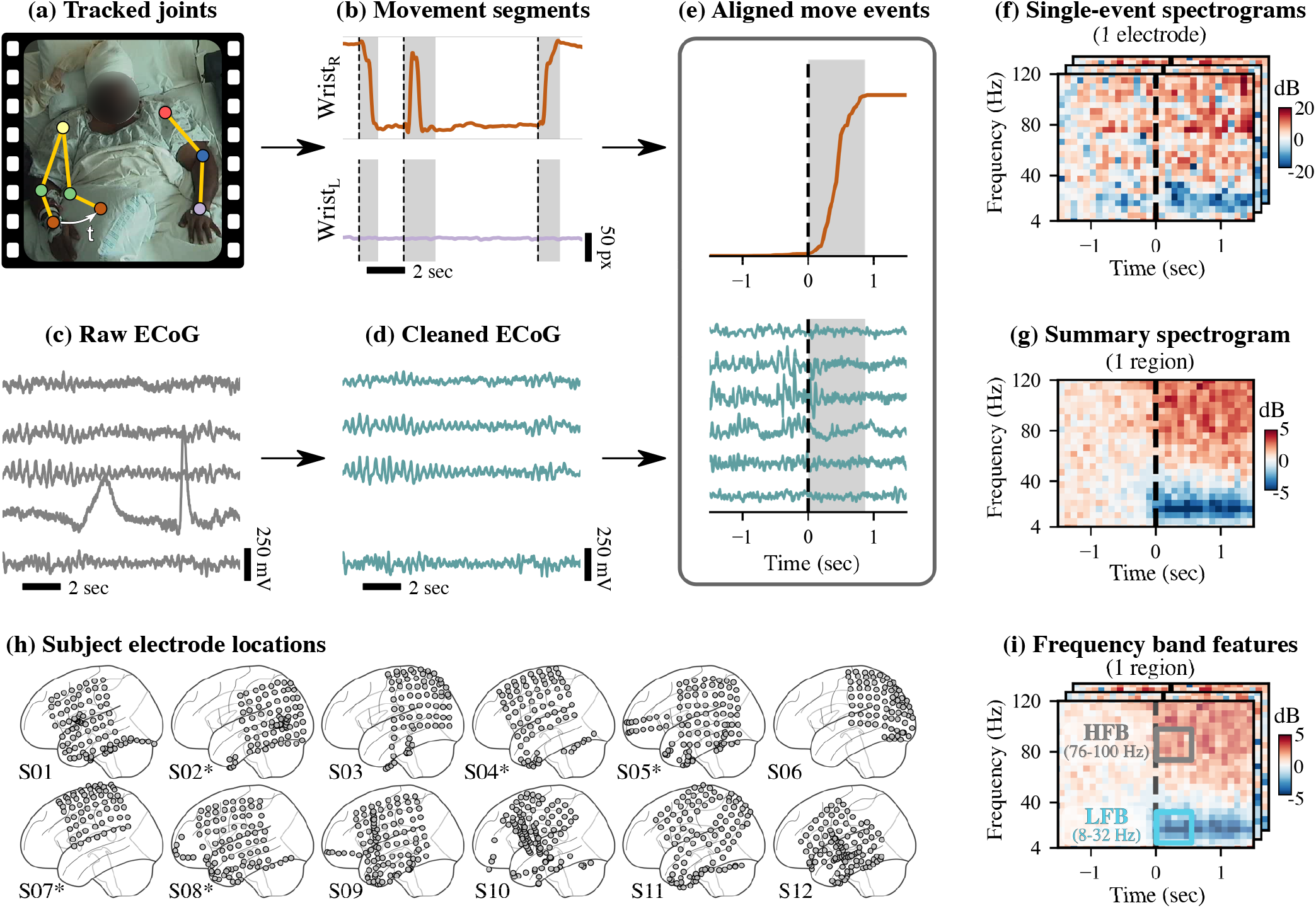
Schematic overview of data processing, analysis, and modeling framework. (a)-(b) Based on continuous video monitoring of each subject, trajectories of the left and right wrists (Wrist_L_ and Wrist_R_ in (b)) were estimated using neural networks [70] and automatically segmented into move (gray) and rest (white) states as shown in (b) [71]. (c)–(d) Raw multielectrode electrocorticography (ECoG) was filtered and re-referenced; bad electrodes (e.g., ones with artifacts) were removed from further analysis. (e) Movement onset events detected from video as shown in (b) were aligned with ECoG data using timestamps. (f) For each move event at each electrode, spectral power was computed and visualized as a log-scaled spectrogram. (g) Summarizing across events and electrodes, we projected the spectral power from electrodes onto 8 cortical regions based on anatomical registration and computed the median power across movement events. (h) Our data included 12 subjects; their electrode placements are shown in MNI coordinates. Five of the subjects had electrodes implanted in their right hemispheres (denoted by asterisks). For consistency of later analyses, we mirrored these electrode locations as shown. (i) To partially explain the event-by-event neural variability in low-frequency (LFB: 8–32 Hz) and high-frequency (HFB: 76–100 Hz) spectral power, we fit multiple linear regression models at each electrode using behavioral features extracted from the videos.

### Behavior during naturalistic movements

The goal of our data processing pipeline was to automate both the identification of wrist movement initiation events and the description of behavioral and environmental features around each event. For each subject, we obtained simultaneously recorded neural activity and movement trajectories immediately before and after the initiation of each movement event (Fig. 1e). Briefly, two-dimensional wrist trajectories were estimated from the video recordings [70] and then segmented into move or rest states [71]. For simplicity of interpretation, we focused on movement initiation events of the wrist contralateral to the ECoG implantation hemisphere, detected during transitions from rest to move states. While we later analyzed ipsilateral wrist behavior to determine if a contralateral wrist movement was bimanual, we did not use the ipsilateral wrist for detecting movement events.

The spontaneous wrist movement events that we identified include a wide variety of upper-limb movement behaviors. Because subjects were sitting in bed, a majority of the movements that we analyzed involved relatively little movement of the shoulders and elbows (see, for example, Supplementary Fig. 1a). Most of the detected movements corresponded to actions such as reaching for a phone, eating, or touching one’s face. We confirmed that our event detection primarily identified contralateral wrist movements, as seen in Supplementary Fig. 1c.

To better assess behavior during wrist movement events, we obtained quantitative values of various movement features and associated environmental variables. We defined a *reach* as the maximum radial displacement of the wrist during the detected movement event, as compared to the wrist position at movement initiation. We extracted 10 behavioral metadata features that quantified the time when each reach began, how the contralateral wrist moved during the reach, whether people were speaking during movement initiation, and how much both wrists moved during each movement [71].

We find that many metadata feature distributions show large within-subject and between-subject variations (Fig. 2). The number of reaches detected across days of recording were fairly consistent, with the exceptions of subjects 04, 05, and 09, who each had one day representing most of the total events. As expected, detected movement events often occurred mostly during waking hours. Reach duration and reach magnitude show minimal inter-subject variability, with most reaches lasting less than 2 seconds and covering fewer than 200 pixels (~67 cm). For reach angle, the distributions tend to be bimodal, with peaks at ±90°, indicating that detected events are biased towards upward and downward reaches, with few side-to-side reaches. Both onset speed and speech ratio distributions vary greatly across subjects, likely reflecting inter-subject differences in the activities performed and the number of people visiting during the detected movement initiations. We also considered a number of features related to coordinated movements with the ipsilateral arm. For bimanual ratio and overlap features, the distributions are skewed towards unimanual movements of the contralateral limb, as expected from Supplementary Fig. 1c, with less skew for subjects 02, 04, and 05. In contrast, the bimanual class categorical feature is primarily skewed towards bimanual movements, indicating that the ipsilateral wrist is often moving, but only a small amount.

**Fig. 2:**
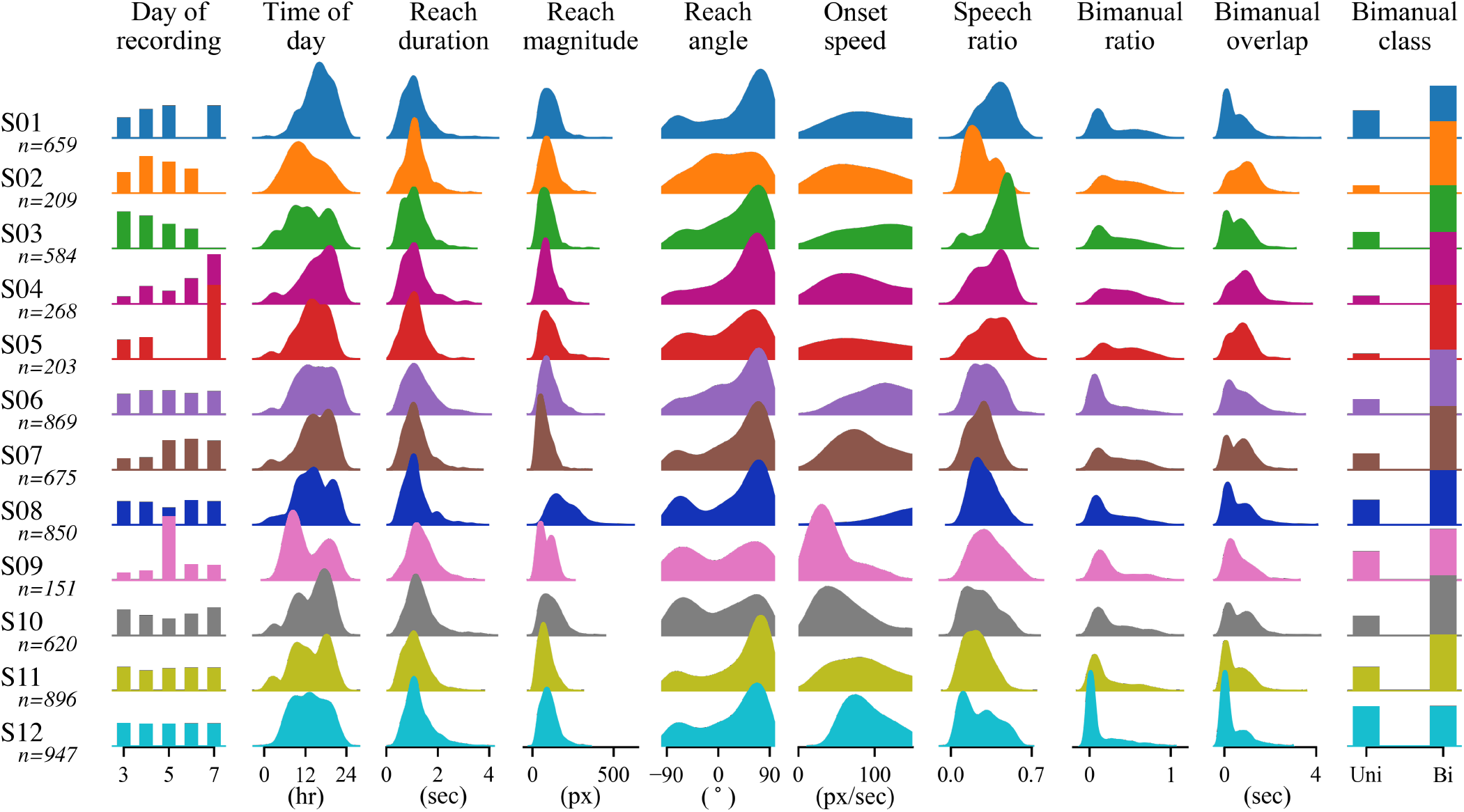
The distribution of extracted behavioral and environmental features show large inter-subject variability. For each subject, features shown include timing (day of recording, time of day), reach parameters (duration, magnitude, angle, onset speed), environment (speech ratio), and bimanual factors (ratio, overlap, and class). The total number of events for each subject was between 151 and 947 (median of 640 across subjects). Each distribution was normalized. These extracted features were used as inputs to the multiple regression models. Note that 3 pixels approximately equal 1 cm.

We also assessed group-level correlations between feature pairs, finding high correlations for 3 reach parameter feature pairs and between all 3 bimanual feature pairs (Supplementary Fig. 2). Reach magnitude positively correlates with reach duration (*r* = 0.26) and onset speed (*r* = 0.56), meaning that reaches tended to cover more distance when they lasted longer or had higher onset speed. Reach duration is also positively correlated with bimanual overlap (*r* = 0.48) due to movements with long durations having more possible overlap time. The high correlations between bimanual features (pairwise Pearson correlation coefficients between overlap v. ratio: *r* = 0.51, class v. ratio: *r* = 0.50, and overlap v. class: *r* = 0.61) indicates that contralateral wrist movements classified as bimanual generally show increased overlap between ipsilateral and contralateral movements and increased ipsilateral amplitude relative to contralateral, as expected.

### Intracortical spectral power during naturalistic movements

We find a consistent set of group-level spectral power patterns, largely localized in fronto-parietal sensorimotor cortical regions. After aligning curated wrist movement events with preprocessed ECoG recordings, we computed time-frequency spectral power at each electrode and then visualized group-level spectral patterns projected onto common regions of interest for all subjects. Generally, we find the expected pattern of low-frequency (~4–30 Hz) spectral power decrease and high-frequency (~50–120 Hz) power increases during movement initiation across multiple cortices (Fig. 3), similar to previous findings during controlled movement experiments [4]. Because ECoG electrode placement varied across subjects, we visualized group-level neural activity by projecting power at every electrode onto 8 common cortical regions of interest [86]: middle frontal, precentral, postcentral, inferior parietal, supramarginal, superior temporal, middle temporal, and inferior temporal. Maximal power deviations primarily occur near movement onset, as expected. Spectral power deviations are largest in magnitude in precentral, postcentral, and inferior parietal regions, which are located in sensorimotor areas of the brain. The middle frontal region also contains strong power fluctuations that could indicate motor planning and possible recruitment of the supplementary motor area. In addition, low-frequency power decreases appear more spatially widespread than high-frequency power increases and are also present in supramarginal and superior temporal regions. As expected, all 3 temporal cortical regions contain minimal movement-related spectral power fluctuations.

**Fig. 3:**
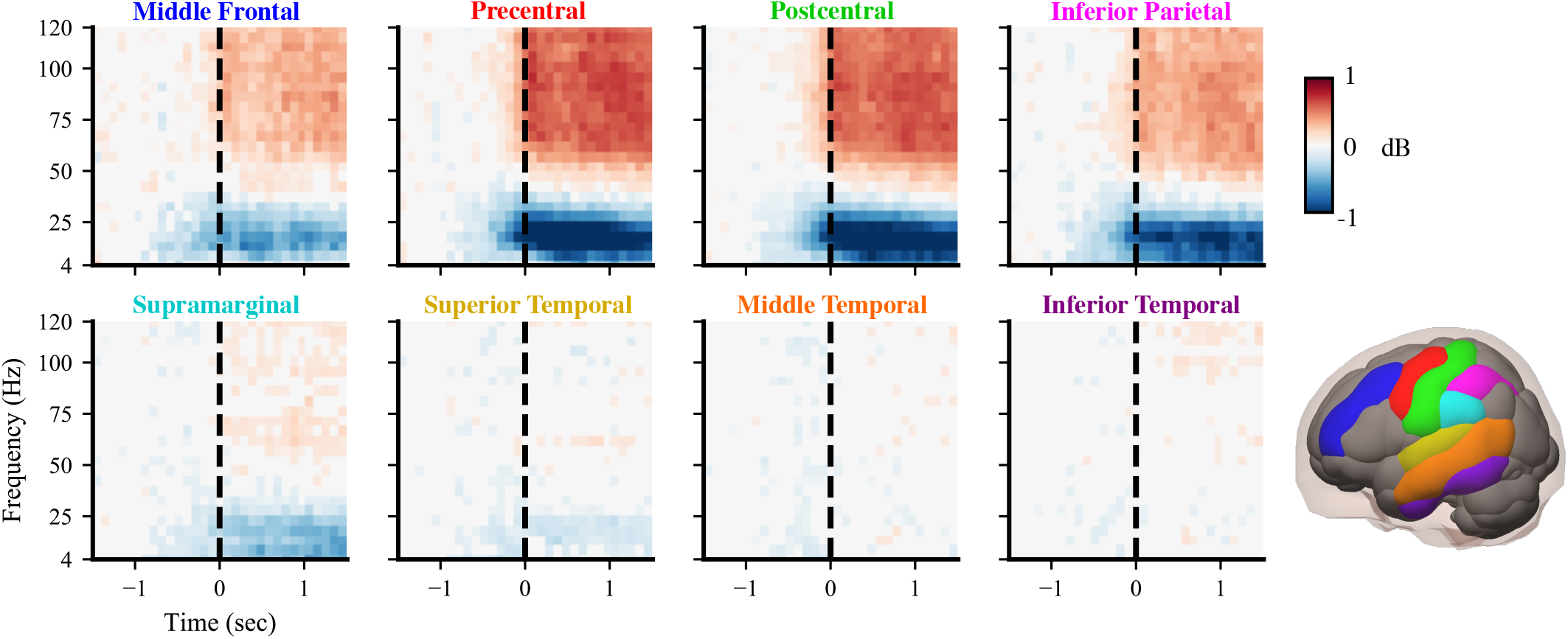
Group-level cortical spectral power changes are consistently localized to sensorimotor regions. Spectrograms show movement event-triggered spectral power patterns for 8 cortical regions (highlighted in lower right) summarized across all 12 subjects. In general, low-frequency (4–30 Hz) power decreases and high-frequency (50–120 Hz) power increases at movement initiation (0 sec), with the largest power fluctuations in fronto-parietal sensorimotor areas. Spectral power was projected based on anatomical registration from electrodes onto 8 regions of interest: middle frontal (blue), precentral (red), postcentral (green), inferior parietal (magenta), supramarginal (cyan), superior temporal (yellow), middle temporal (orange), and inferior temporal (purple). We subtracted the baseline power of 1.5–1 seconds before movement initiation. Non-significant differences from baseline power were set to 0 (*p* > 0.05).

Despite consistent group-level spectral power patterns across cortical regions, we identified considerable spectral power variability across subjects (Supplementary Fig. 3). For instance, the precentral region shows the same low/high-frequency power pattern for each subject (Fig. 4), but the amplitudes and frequency bands of maximal power deviation differ widely across subjects. Subjects 03, 06, 07, 08, and 11 show increased power at high frequencies up to 120 Hz, while subjects 09 and 12 have increased power primarily between 60–80 Hz. For subjects 04 and 08, low-frequency power decreases occur across narrower frequency bands compared to the other subjects. Besides arising from inter-subject differences in neural anatomy and connectivity, these spectral power variations may reflect variability in daily activities, electrode placement, medication, and seizure foci among subjects [87, 88]. Spectral power plots for the 7 other regions of interest are shown in Supplementary Figs. 4– 10.

**Fig. 4:**
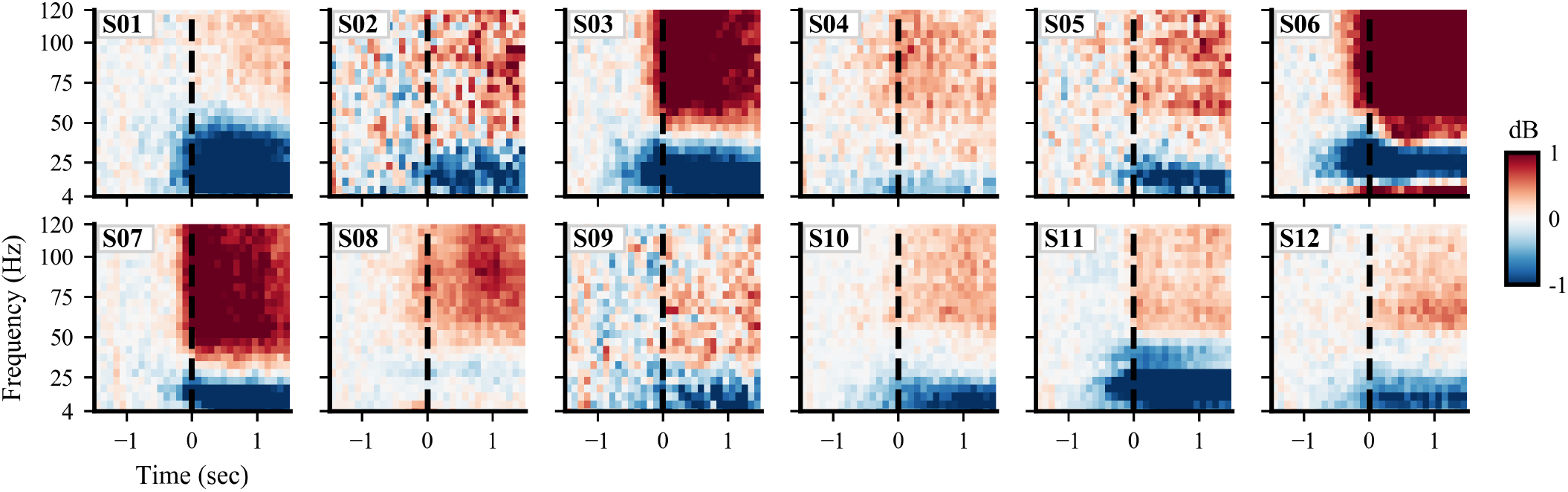
Spectral power patterns in the precentral region vary considerably across subjects. While some subjects show spectral power patterns similar to the group-level results in Fig. 3, many deviate substantially from the group average pattern in both magnitude and frequency bands. The colormap indicates differences in spectral power relative to baseline 1.5–1 seconds before movement initiation (no statistical masking is used).

In addition to inter-subject neural variability, we also identified notable changes in movement-related neural activity across recording days for several subjects (Fig. 5). We analyzed spectral power in the precentral region averaged over the half second following movement onset and split into low-frequency (LFB: 8–32Hz) and high-frequency (HFB: 76–100Hz) bands, similar to previous research [4]. For both frequency bands, spectral power significantly differed across subjects (p<0.001 for both bands, Kruskal-Wallis test). We also found a significant effect of recording day in LFB for subjects 03 (p<0.001, Kruskal-Wallis test), 05 (p=0.013), 06 (p<0.001), 07 (p=0.041), 08 (p=0.013), and 11 (p<0.001) as well as in HFB for subjects 03 (p=0.002), 04 (p<0.001), 07 (p<0.001), 08 (p=0.004), and 10 (p=0.006). Surprisingly, these significant recording day effects appear for several subjects despite baseline-subtracting spectral power features. Yet, baseline subtraction does substantially reduce neural variability across recording days (Supplementary Fig. 11), as expected.

**Fig. 5:**
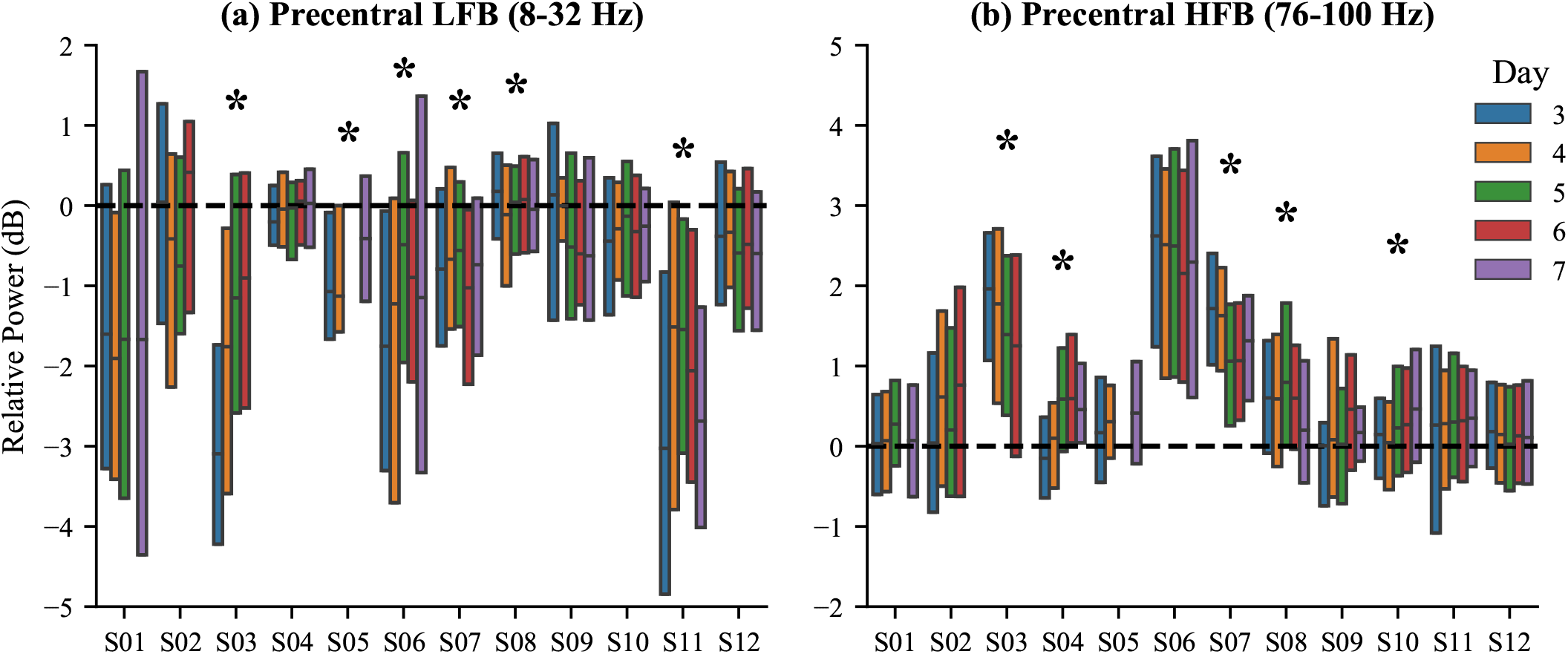
Precentral banded spectral power varies considerably across subjects and recording days. (a) Low-frequency (LFB: 8–32 Hz) and (b) high-frequency (HFB: 76–100 Hz) spectral power in the precentral region was averaged over the first half-second after movement onset. Boxplots show spectral power variability across events for every subject, separated by recording day. For each subject, a significant recording day effect on spectral power is denoted by an asterisk (p<0.05, Kruskal-Wallis test).

### Modeling single-event spectral power with behavioral features

We developed a robust multiple variable linear regression model to explain single-event spectral power at each intracranial electrode using our 10 behavioral metadata features (Fig. 6a). For each electrode, we modelled LFB and HFB spectral power averaged over the half second following movement onset (Fig. 1i). For every model, behavioral features were pruned independently using forward selection to avoid overfit-ting. We assessed model performance on randomly withheld movement events by computing an *R*^2^ score (referred to as the full model *R*^2^). To assess the contributions of each individual feature, we shuffled that feature’s training labels, fit a new linear model, and subtracted this model’s *R*^2^ on withheld data from the full model *R*^2^ to obtain an estimate of feature importance. Higher Δ*R*^2^ values indicate features that explain more variance.

**Fig. 6:**
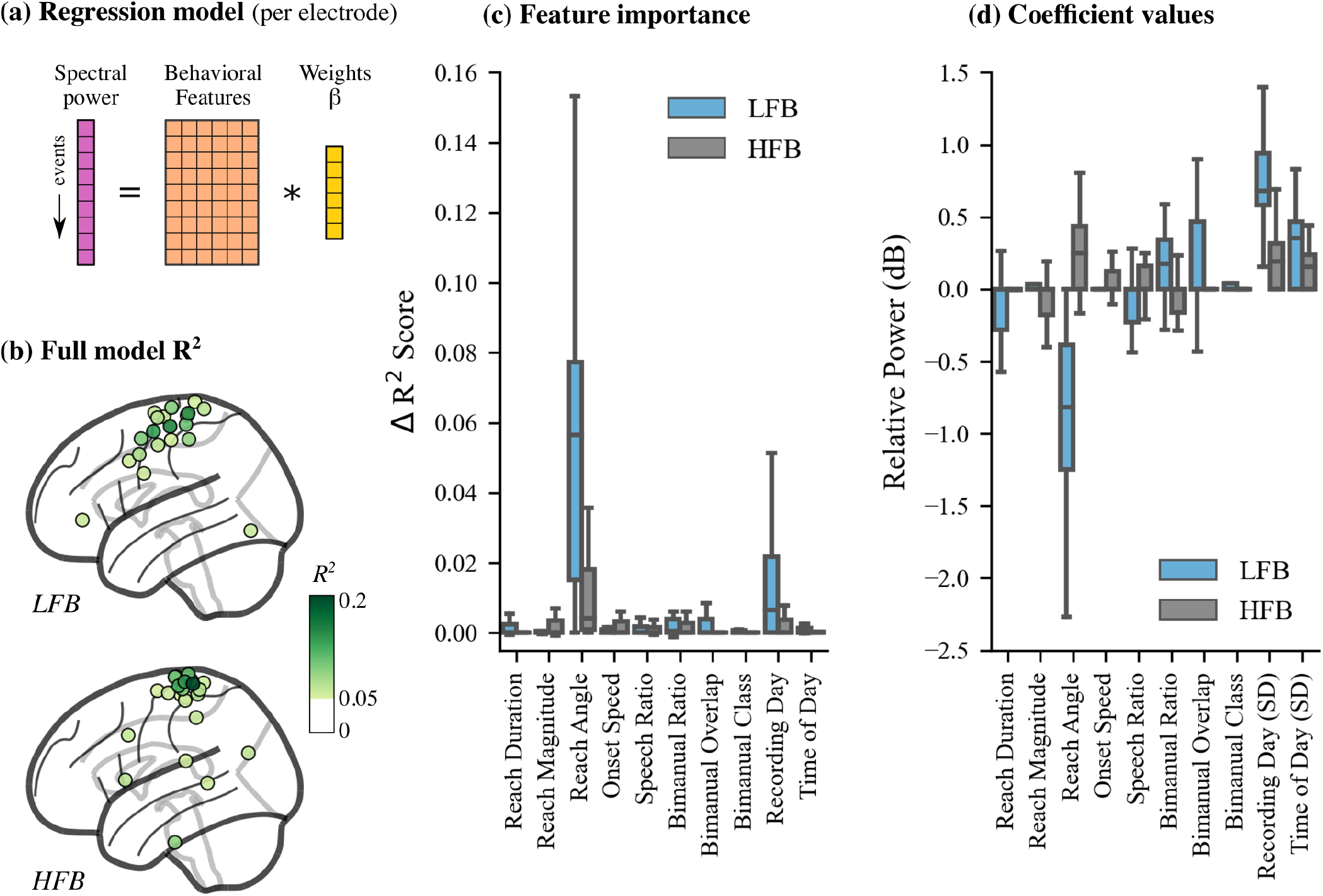
Event-by-event multiple regression models explain changes in neural spectral power using extracted behavioral and environmental features. (a) We fit multiple linear regression models at each electrode using behavioral features extracted from the videos (orange) and frequency-banded spectral power (magenta). (b) Models with the largest *R*^2^ scores on withheld data were primarily located in sensorimotor areas. (c) Reach angle and recording day were the most explanatory model features, especially when regressing low-frequency spectral power. (d) Regression coefficients indicate that upward reaches enhance the average spectral power pattern observed. Recording day and time of day both have large standard deviations (SD) across one-hot encoded variable coefficients, highlighting the effects of long-term temporal variability Only models with *R*^2^ > 0.05 on withheld data are shown for (b)–(d).

For both frequency bands, intracortical activity variability is best explained by our models fit to electrodes located in fronto-parietal sensorimotor areas (Fig. 6b and Supplementary Fig. 12). This finding matches well with the spatial distribution of spectral power (see Fig. 3). However, all *R*^2^ scores are at most 0.25, indicating that even the best models cannot explain more than 75% of the variance in the withheld data. Among individual features, we find that reach angle and day of recording are the most informative (Fig. 6c). Reach angle is also the most often retained feature following forward selection in sensorimotor regions (Supplementary Table 1), indicating its importance for modelling neural activity during movement onset. In addition, both reach angle and day of recording have the largest coefficient magnitudes among behavioral features in the regression models (Fig. 6c). The coefficients for reach angle indicate that upward reaches are associated with decreased low frequency power and increased high frequency power compared to the average response. In other words, upward reaches tend to increase the magnitude of the spectral power pattern seen. The SD of the coefficients corresponding to recording day are surprisingly large, indicating that baseline-subtracted neural responses still vary across long time scales usually not captured in short, controlled experiments. This observation highlights the importance of properly accounting for long-term temporal effects when understanding and decoding neural recordings.

## Discussion

Our results demonstrate that electrocortical correlates of naturalistic arm movements in humans corroborate findings from controlled experiments on average, as we had hypothesized. However, we found high behavioral and neural variability during naturalistic movements across participants and recording days. Using multiple regression modelling, we were able to partially explain this event-by-event electrocortical variability using behavioral metadata features extracted from video recordings. In general, we find that results from controlled upper-limb reaching tasks do generalize to naturalistic movements on average, but naturalistic movements involve considerable event-by-event neural variability that cannot be fully explained by simple behavioral and environmental measures.

Across subjects, we observe an average decrease in low-frequency band cortical power and increase in high-frequency band cortical power during naturalistic upper-limb movement initiation, consistent with previous controlled studies [4, 64]. Decreases in low-frequency power are thought to reflect changes in the current neural state if a new or unexpected event occurs [89]. In our study, the neural state can be disrupted during movement initiation by a variety of factors, such as increased attention or prediction error once the arm is in motion. In contrast, high-frequency power increases may indicate active sensorimotor processing [90–94]. Low-frequency and high-frequency power changes are thought to represent two separate processes [95, 96], which could explain the difference seen in the spatial spread of cortical power changes between low and high frequencies. In our study, the frequency bands of the maximum spectral power responses do differ across subjects, suggesting that the processes underlying the low and high frequency bands vary across subjects. This inter-subject variability reinforces the importance of assessing both subject-specific neural responses and group-level activity.

Despite showing the expected cortical pattern on average, naturalistic reaches exhibited notable behavioral and neural variability across subjects and recording days. This high variability may reflect variations in sensory input and movement constraints due to different types of behaviors [97]. Categorizing such behaviors during reaching would be challenging, however, due to many possible neurally-relevant behavior types and a lack of objective measures that can properly discriminate such behaviors without user-defined labels [98]. The neural variation seen across recording days could be caused by several factors, including changes in medication, seizure frequency, and alertness while recovering from ECoG implantation surgery. Similar long-term, inter-day variability has been observed in previous EEG and ECoG studies [99–101]. It is also worth noting that these day-to-day changes in ECoG spectral power are small in magnitude (±1–2 dB) relative to spectral power without any baseline subtracted (~10–50 dB). Furthermore, recent research suggests that despite long-term neural recording variability, low-dimensional representations of this activity remain stable over long periods of time [102].

During regression modelling, we find that our models only explained at most 25% of the variability; this measure is low, but not unusual given the single-event noise in the electrocortical signal [103]. Furthermore, some of this variability may be explained by other movement behaviors beyond what we quantified using our pose tracking methodology [104]. These low scores may also reflect the simplicity of our linear models. While studies have shown evidence of nonlinear relationships between electrocortical activity and behavior [28, 105], we chose linear regression models because they provide easily interpretable results and allow straightforward assessments of individual feature contributions.

Our regression model identified vertical reach angle and day of recording as the most explanatory features. The importance of vertical reach angle makes sense because upward reaches require more effort and activate different muscles than downward reaches. In addition, population neural activity has been shown to robustly encode reach direction [106, 107]. We did not include a reach angle feature sensitive to horizontal movements because reach angle distributions were skewed towards vertical angles at ±90°, as seen in Fig. 2. The day of recording feature was also found to explain some of the neural variance captured by regression modelling. This finding is sensible given the significant inter-day neural variability seen for several subjects.

Our study has several important limitations. Because we are performing human ECoG research, we are studying subjects who have epilepsy and are recovering from electrode implantation surgery, which may introduce confounding effects due to medication and seizure location. To address this issue, we ignored data from the first 2 days post-surgery, removed electrodes with abnormal activity, and assessed movements across multiple days to avoid single-day bias. Another limitation is that the clinical video monitoring system includes only one camera, whose view can be obstructed by people and various objects throughout the day. We minimized obstruction effects by selecting movement events with high confidence scores in the event detection algorithm [71] and manually reviewing all detected events to check if they were actual movements and not false positives, but using multiple cameras would extend body tracking to 3D in future studies. Additionally, the video camera was positioned by the clinical staff and was vertically rotated away by the clinical staff during private times, meaning that the video view changed slightly throughout the day. We minimized the effect of such rotations on our behavioral features by excluding camera rotation events and using movement features that were relative to the start of each reach.

Our results underline the importance of studying naturalistic movements and understanding neural variability across multiple days. Our approach leverages pre-existing clinical setups and could be extended to other movements and behaviors, such as grasping objects, sleep/wake transitions, and conversing with others. More broadly, our results have implications for developing novel brain-computer interfaces that can decode neural data across subjects in natural environments. For instance, movement data from many subjects cold be combined to train decoders that generalize to new subjects with minimal re-training and are robust to a richer set of behavioral and environmental contexts. By publicly releasing our curated dataset, we hope to spur further research that enhances our understanding of naturalistic behavior and informs the development of nextgeneration brain-computer interfaces.

## Methods

### Subject information

We analyzed opportunistic clinical recordings from 12 subjects (8 males, 4 females) during their clinical epilepsy monitoring (conducted at Harborview Medical Center in Seattle, WA). Subjects were 29.4±7.9 years old at the time of recording (mean±SD). Our study was approved by the University of Washington Institutional Review Board for the protection of human subjects. All subjects provided written informed consent.

We selected subjects who had ECoG electrode coverage near primary motor cortex, with either one 8×8 or two 4×8 electrode grids placed subdurally on the cortical surface. Additional electrodes were implanted on the cortical surface for some subjects, resulting in 87.0± 12.9 total surface electrodes per subject (mean±SD). In addition, five subjects had 23.2±12.1 intracortical depth electrodes (mean±SD). Electrodes were implanted primarily within one hemisphere for each subject (5 right hemisphere, 7 left hemisphere). Single-subject electrode placement and recording duration information are given in Supplementary Table 2.

### Data collection

Subjects underwent 24-hour clinical monitoring, involving semi-continuous ECoG and audio/video recordings over 7.4±2.2 days per subject (mean±SD). Some breaks occurred throughout monitoring (on average, 8.3±3.2 total breaks per subject, each lasting 1.9±2.4 hours [mean±SD]). For all subjects, we restricted our analysis to days 3–7 following the electrode implantation surgery, in order to exclude potentially anomalous neural and behavioral activity immediately following electrode implantation surgery. For several subjects, some days were excluded due to corrupted or missing data files, as noted in Supplementary Table 2. During clinical monitoring, subjects were observed during a variety of typical everyday activities, such as eating, sleeping, watching television, and socializing while confined to a hospital bed. ECoG and video were initially sampled at 1000 Hz and 30 frames per second, respectively. Fig. 1 shows an example of the clinical monitoring setup, along with our data processing pipeline.

### ECoG data processing

We processed the raw ECoG data using custom MNE-Python scripts [108]. First, we removed DC drift by subtracting the median voltage of each electrode. Widespread, high-amplitude artifacts were then identified by abnormally high electrode-averaged absolute voltage (> 50 interquartile range [IQR]). We set these artifacts to 0, along with all data within 2 seconds of each identified artifact. Removing such high-amplitude discontinuities minimizes subsequent filtering artifacts due to large, abrupt changes in the signal [109].

With data discontinuities removed, we band-pass filtered the data (1–200 Hz) and notch filtered to minimize line noise at 60 Hz and its harmonics. The data were then resampled to 500 Hz and re-referenced to the common median for each grid, strip, or depth electrode group. Electrodes with bad data were identified based on abnormal standard deviation (> 5 IQR) or kurtosis (> 10 IQR) compared to the median value across channels. This process resulted in the removal of 4.9±4.9 surface electrodes per subject and 1.0±1.4 depth electrodes for each of the 5 subjects with depth electrodes.

Electrode positions were localized using the Fieldtrip toolbox in Matlab [110, 111] to enable multi-subject analyses. This process involved co-registering preoperative MRI and postoperative CT scans, manually selecting electrodes in 3D space, and warping electrode positions into MNI space.

### Movement event identification and pruning

We performed markerless pose estimation on the raw video footage separately for each subject to determine wrist positions (Fig. 1a). First, for each subject, we manually annotated 1000 random video frames with the 2D positions of 9 keypoints: the subject’s nose, ears, wrist, elbows, and shoulders (https://tinyurl.com/human-annotation-tool). Video frames were randomly selected across all days, with preference given to frames during active, daytime periods. These manually annotated frames were used to train a separate neural network model for each subject using DeepLabCut [70]. Each model was then applied to every video for that subject to generate estimated wrist trajectories.

Movement states were identified by applying a first-order autoregressive hidden semi-Markov model to each wrist trajectory. This state segmentation model classified the wrist trajectory into either a move or rest state. For this study, we focused on movements of the wrist contralateral to the implanted hemisphere. Contralateral wrist states were then discretized, and movement initiation events were identified at state transitions where 0.5 seconds of rest states are followed by 0.5 seconds of move states. See Singh et al. [71] for further methodological details.

After identifying movement initiation events, we coarsely labeled the video data manually (~3 minutes resolution) and excluded arm movements during sleep, unrelated experiments, and private times (as specified in our IRB protocol). In addition, we only retained movement events where (1) movement durations were between 0.5–4 seconds, (2) the confidence scores from DeepLabCut were > 0.4, indicating minimal marker occlusion, and (3) wrist movements followed a parabolic trajectory, as determined by a quadratic fit to the wrist’s radial movement (*R*^2^ > 0.6). We found that this quadratic fit criteria eliminated many outliers with complex movement trajectories and improved the interpretability of our subsequent analyses. For each day of recording, we selected up to 200 events with the highest movement onset velocities. Finally, all movement initiation events were visually inspected, and events with occlusions or false positive movements were removed (17.8%±9.9% of events [mean±SD]).

### ECoG-event synchronization and segmentation

We used timestamps accompanying clinical recordings to synchronize movement initiation events with ECoG recordings and generated 10-second ECoG segments centered around each event. ECoG segments with missing data and large artifacts, such as line noise, were removed by computing log-transformed spectral power density for each segment and discarding segments with power below 0 dB or with abnormally high power at 115–125 Hz (>3 SD) compared to all segments. With these bad ECoG segments removed, we computed log-transformed, time-frequency spectral power using Morlet wavelets [112]. Power at each segment was then baseline-subtracted, using a baseline defined as 1.5–1 seconds before each movement initiation event.

### Projecting power into regions of interest

Because electrode placement was clinically motivated and varied greatly across subjects, we projected the spectral power computed at every electrode into common regions of interest defined by the AAL atlas [113]. Prior to projection, in order to combine all subjects, all right hemisphere electrode positions were flipped into the left hemisphere. Using EEGLAB and Matlab, we mapped from electrodes to small, predefined brain regions by positioning a three-dimensional Gaussian (2 cm full-width at half-maximum) centered at each electrode position and calculating the Gaussian’s value at each small region [86, 114, 115]. The values across small regions were combined based on the AAL region boundaries, providing a mapping between each electrode and AAL region based on radial distance. We performed this projection procedure separately for each subject.

By summing the weights from these mappings across electrodes, we estimated the electrode density for each AAL region. We retained regions with an average electrode density > 3 across subjects, resulting in 8 regions of interest (ROIs): middle frontal, precentral, postcentral, inferior parietal, supramarginal, superior temporal, middle temporal, and inferior temporal (Fig. 3). These 8 ROIs represent where most of the electrodes were located across subjects. We then normalized the weights for each ROI so that they summed to 1. These normalized weights were used to perform a weighted average of electrode-level spectral power for every ECoG segment, generating a spectral power estimate at each region of interest.

After projecting single-event spectral power onto regions of interest, we computed the median value across events separately for each subject and region. We then averaged the event-median spectral power across subjects to obtain group-level estimates for each region of interest. To mask spectral power patterns that were not significant, group-level spectral power for every frequency bin within each region of interest was then compared to a 2000-permutation bootstrap distribution generated from baseline time points. Non-significant differences from each bootstrap distribution were set to 0 (*p* > 0.05, two-sided bootstrap statistics, false discovery rate correction [116]).

### Single-event behavioral metadata features

We extracted multiple behavioral and environmental metadata features that quantify variations in movement parameters and environmental contexts. These features were later used as input variables for regression models of inter-event spectral power and can be divided into 4 categories.

#### 1) Timing features

Day of recording and time of day for each movement initiation event are used to capture long-term variations in the neural response.

#### 2) Reach movement features

To quantify differences in the detected movements, we defined a reach as the maximum radial displacement of the wrist marker during the detected move state compared to its position at each movement initiation event. These features included the duration and magnitude of each reach. We also computed the 2D reach angle and transformed angles at 90–270° to range from 90° to −90°, respectively. This transformation made the reach angle sensitive to vertical reach variations, with 90° for upward reaches and −90° for downward reaches. We also computed wrist marker radial speed during movement onset. Note that these movement features were based on the location of the video camera, which varied slightly across subjects and recording days.

#### 3) Environmental feature

Based on results from the literature [81, 82], we were motivated to consider how environmental factors affect electrocortical power. Here, we examined the environmental factor of people talking during movement initiation. First, we cleaned the recorded audio signal using spectral noise gating (https://www.audacityteam.org), which performed 40 dB reduction on audio signal components that were similar to a selected noise period during rest. We then used the short-time Fourier transform to compute the spectral power from 370–900 Hz as a proxy for speech [117]. This power was divided by the total power at each time point, producing a ratio that is robust to broadband changes in the audio signal caused by noise. This speech ratio was smoothed using a 1st-order low-pass filter with 4.2 mHz cutoff to minimize the effects of transient changes in power due to noise. We then averaged this ratio from −1 to 1 seconds around each movement initiation event, generating a speech ratio feature that ranges from 0.0 to 1.0.

#### 4) Bimanual reach features

While movement initiation event selection was based solely on contralateral wrist movement, the ipsilateral wrist can still move and may affect the electrocortical response. We quantified the relative magnitude of ipsilateral wrist movement by computing the ratio of the ipsilateral wrist reach magnitude to the sum of ipsilateral and contralateral reach magnitudes. In addition, we computed the temporal overlap between contralateral and ipsilateral move states over the duration of the entire contralateral wrist movement. Finally, we computed a binary feature that classified movements as either unimanual or bimanual based on the amount of temporal lag between contralateral and ipsilateral wrist movement onset. This feature was bimanual if a sequence of 4 consecutive move states of the ipsilateral wrist began either 1 second before contralateral wrist movement initiation or anytime during the contralateral wrist move state.

### Single-event spectral power linear regression

Using the 10 extracted behavioral features as independent variables, we fit a separate linear regression model to the spectral power at every electrode. While projecting onto cortical regions provided a useful visualization, we found that fitting regression models using projected power resulted in very poor model fits, likely due to electrodes with maximal power responses overlapping multiple regions and differing across subjects. All features were standardized prior to regression, with reach duration and reach magnitude features also being log-transformed. We categorized the two timing features using one-hot encoding based on day of recording and three 8-hour segments (12am– 8am, 8am–4pm, 4pm–12am) for time of day because we do not expect linear long-term power changes within and across days. For the dependent variable, we averaged spectral power over the first half second of movement onset for low-frequency (8–32 Hz) and high-frequency (76–100 Hz) bands. We then randomly selected 90% of each subject’s total contralateral arm movement events as training data, while withholding the remaining 10% for testing model generalizability. For each model, we independently pruned input features using forward selection, retaining features that improved adjusted *R*^2^ for an ordinary least squares fit. This procedure helped minimize overfitting due to too many independent variables.

For training, we applied a multiple linear regression model for event-by-event spectral power patterns (shown schematically in Fig. 1i) defined as:

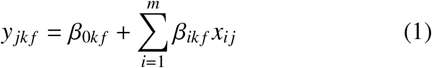

where *y_jkf_* is the spectral power for movement event *j* at electrode *k* averaged over frequency band *f*, during the first half second of movement initiation; *x_ij_* is feature *i* at event *j*, and *β_ikf_* is the coefficient for feature *i* at electrode *k* and frequency band *f* (*β_0kf_* is the intercept term). We minimized the Huber norm during model fitting to improve model robustness to outliers [118].

After training, we performed model validation by computing the *R*^2^ on withheld data, referred to as the full model *R*_2_. We also assessed the contribution of each behavioral feature independently by shuffling one feature, fitting a new model, and computing the *R*^2^ on the unshuffled, withheld data. This new *R*^2^ was subtracted from the full model *R*^2^ to obtain Δ*R*^2^ as an estimate of that feature’s importance. We repeated this shuffling process and computation of Δ*R*^2^ across all model features.

We computed independent regression models using forward selection, along with *R*^2^ and Δ*R*^2^ scores, over all electrodes and for both low and high frequency bands. To minimize bias in our selection of training and testing data, we performed 200 random, independent train/test splits for every regression model, averaging the full model *R*_2_, Δ*R*^2^, and coefficients across all splits. We balanced days of recording within each train and test set.

## Data availability

Our curated dataset is publicly available without restriction, other than citation, through Figshare at https://figshare.com/projects/Behavioral_and_neural_variability_of_naturalistic_arm_movements/78666. This public dataset contains synchronized neural and behavioral data that can be used to generate Fig. 2–6.

## Code availability

Our data analysis code is publicly available without restriction, other than citation, on Github at https://github.com/BruntonUWBio/naturalistic_arm_movements_ecog. The code in this repository can be used in conjunction with our published dataset to reproduce all main findings and figures from our study. We ask that future studies building on our published data and code cite both this paper and [71].

## Acknowledgements

We thank John So for his extensive manual annotations of the video data; Kameron D. Harris, Nile Wilson, Kurt Weaver, and Leigh Weber for discussions and help with data collection and analysis; neurosurgeons Jeffrey G. Ojemann and Andrew Ko, as well as the excellent staff at Harborview Hospital Neurosurgery department, for their care of the patients during their monitoring. This research was supported by funding from the Defense Advanced Research Projects Agency (FA8750-18-2-0259), the National Science Foundation (1630178 and EEC-1028725), the Sloan Foundation, and the Washington Research Foundation.

## Author contributions

NXRW, RPNR, and BWB conceived of the study. SMP and SHS performed the data analysis. SMP, SHS, RPNR, and BWB interpreted the results. SMP and BWB wrote the manuscript. SMP, SHS, NXRW, RPNR, and BWB edited the final manuscript. RPNR and BWB acquired funding for the project.

## References

[1] Sober, S. J., Sponberg, S., Nemenman, I. & Ting, L. H. Millisecond spike timing codes for motor control. Trends in Neurosciences 41, 644–648 (2018).

[2] Kalaska, J. F. From Intention to Action: Motor Cortex and the Control of Reaching Movements, 139–178 (Springer US, Boston, MA, 2009).

[3] Truccolo, W., Friehs, G. M., Donoghue, J. P. & Hochberg, L. R. Primary motor cortex tuning to intended movement kinematics in humans with tetraplegia. The Journal of Neuroscience 28, 1163 (2008).

[4] Miller, K. J. et al. Spectral changes in cortical surface potentials during motor movement. Journal of Neuroscience 27, 2424–2432 (2007).

[5] Nakanishi, Y. et al. Prediction of three-dimensional arm trajectories based on ecog signals recorded from human sensorimotor cortex. PloS one 8, e72085 (2013).

[6] Wang, Z. et al. Decoding onset and direction of movements using electrocorticographic (ecog) signals in humans. Frontiers in neuroengineering 5, 15 (2012).

[7] Schalk, G. et al. Two-dimensional movement control using electrocorticographic signals in humans. Journal of neural engineering 5, 75 (2008).

[8] Georgopoulos, A. P., Merchant, H., Naselaris, T. & Amirikian, B. Mapping of the preferred direction in the motor cortex. Proceedings of the National Academy of Sciences 104, 11068 (2007).

[9] Leuthardt, E. C., Schalk, G., Wolpaw, J. R., Ojemann, J. G. & Moran, D. W. A brain–computer interface using electrocorticographic signals in humans. Journal of neural engineering 1, 63 (2004).

[10] Umeda, T., Koizumi, M., Katakai, Y., Saito, R. & Seki, K. Decoding of muscle activity from the sensorimotor cortex in freely behaving monkeys. NeuroImage 197, 512–526 (2019).

[11] Fried, I., Haggard, P., He, B. J. & Schurger, A. Volition and action in the human brain: Processes, pathologies, and reasons. The Journal of neuroscience: the official journal of the Society for Neuroscience 37, 10842–10847 (2017).

[12] Kornhuber, H. H. & Deecke, L. Brain potential changes in voluntary and passive movements in humans: readiness potential and reafferent potentials. Pflügers Archiv - European Journal of Physiology 468, 1115–1124 (2016).

[13] Jackson, A., Mavoori, J. & Fetz, E. E. Correlations between the same motor cortex cells and arm muscles during a trained task, free behavior, and natural sleep in the macaque monkey. Journal of Neurophysiology 97, 360–374 (2007).

[14] Lee, I. H. & Assad, J. A. Putaminal activity for simple reactions or self-timed movements. Journal of Neurophysiology 89, 2528–2537 (2003).

[15] Romo, R. & Schultz, W. Neuronal activity preceding self-initiated or externally timed arm movements in area 6 of monkey cortex. Experimental Brain Research 67, 656–662 (1987).

[16] Nastase, S. A., Goldstein, A. & Hasson, U. Keep it real: rethinking the primacy of experimental control in cognitive neuroscience (2020).

[17] Dastjerdi, M., Ozker, M., Foster, B. L., Rangarajan, V. & Parvizi, J. Numerical processing in the human parietal cortex during experimental and natural conditions. Nature Communications 4, 2528 (2013).

[18] Wilson, N. R. et al. Cortical topography of error-related high-frequency potentials during erroneous control in a continuous control braincomputer interface. Frontiers in Neuroscience 13, 502 (2019).

[19] Omedes, J., Schwarz, A., Müller-Putz, G. R. & Montesano, L. Factors that affect error potentials during a grasping task: toward a hybrid natural movement decoding bci. Journal of Neural Engineering 15, 046023 (2018).

[20] Gilja, V. et al. Challenges and opportunities for nextgeneration intracortically based neural prostheses. IEEE Transactions on Biomedical Engineering 58, 1891–1899 (2011).

[21] Schalk, G. et al. Two-dimensional movement control using electrocorticographic signals in humans. Journal of Neural Engineering 5, 75–84 (2008).

[22] Taylor, D. M., Tillery, S. I. H. & Schwartz, A. B. Direct cortical control of 3d neuroprosthetic devices. Science 296, 1829–1832. (2002).

[23] Kubota, K. & Niki, H. Prefrontal cortical unit activity and delayed alternation performance in monkeys. Journal of Neurophysiology 34, 337–347 (1971).

[24] Fetz, E. E. Operant conditioning of cortical unit activity. Science 163, 955 (1969).

[25] Bizzi, E. Discharge of frontal eye field neurons during saccadic and following eye movements in unanesthetized monkeys. Experimental Brain Research 6, 69–80 (1968).

[26] Evarts, E. V. A technique for recording activity of subcortical neurons in moving animals. Electroencephalography and Clinical Neurophysiology 24, 83–86 (1968).

[27] Evarts, E. V. Relation of pyramidal tract activity to force exerted during voluntary movement. Journal of Neurophysiology 31, 14–27 (1968).

[28] David, S. V., Vinje, W. E. & Gallant, J. L. Natural stimulus statistics alter the receptive field structure of v1 neurons. Journal of Neuroscience 24, 6991–7006. (2004).

[29] Chandrasekaran, C., Trubanova, A., Stillittano, S., Caplier, A. & Ghazanfar, A. A. The natural statistics of audiovisual speech. PLoS computational biology 5, e1000436–e1000436 (2009).

[30] Chandrasekaran, C., Turesson, H. K., Brown, C. H. & Ghazanfar, A. A. The influence of natural scene dynamics on auditory cortical activity. The Journal of Neuroscience 30, 13919 (2010).

[31] Silbert, L. J., Honey, C. J., Simony, E., Poeppel, D. & Hasson, U. Coupled neural systems underlie the production and comprehension of naturalistic narrative speech. Proceedings of the National Academy of Sciences 111, E4687–E4696. (2014).

[32] Huth, A. G., de Heer, W. A., Griffiths, T. L., Theunissen, F. E. & Gallant, J. L. Natural speech reveals the semantic maps that tile human cerebral cortex. Nature 532, 453–458 (2016).

[33] Zuo, X. et al. Temporal integration of narrative information in a hippocampal amnesic patient (2019).

[34] Machens, C. K., Wehr, M. S. & Zador, A. M. Linearity of cortical receptive fields measured with natural sounds. Journal of Neuroscience 24, 1089–1100. (2004).

[35] Chang, L. & Tsao, D. Y. The code for facial identity in the primate brain. Cell 169, 1013–1028.e14 (2017).

[36] Scanlon, J. E. M., Townsend, K. A., Cormier, D. L., Kuziek, J. W. P. & Mathewson, K. E. Taking off the training wheels: Measuring auditory p3 during outdoor cycling using an active wet eeg system. Brain Research 1716, 50–61 (2019).

[37] Nordin, A. D., Hairston, W. D. & Ferris, D. P. Human electrocortical dynamics while stepping over obstacles. Scientific Reports 9, 4693 (2019).

[38] Ladouce, S., Donaldson, D. I., Dudchenko, P. A. & Ietswaart, M. Understanding minds in real-world environments: Toward a mobile cognition approach. Frontiers in human neuroscience 10, 694–694 (2017).

[39] Zink, R., Hunyadi, B., Huffel, S. V. & Vos, M. D. Mobile eeg on the bike: disentangling attentional and physical contributions to auditory attention tasks. Journal of Neural Engineering 13, 046017 (2016).

[40] Gwin, J. T., Gramann, K., Makeig, S. & Ferris, D. P. Electrocortical activity is coupled to gait cycle phase during treadmill walking. NeuroImage 54, 1289–1296 (2011).

[41] Tsitsiklis, M. et al. Single-neuron representations of spatial targets in humans. Current Biology 30, 245–253.e4 (2020).

[42] Maidenbaum, S., Miller, J., Stein, J. M. & Jacobs, J. Grid-like hexadirectional modulation of human entorhinal theta oscillations. Proceedings of the National Academy of Sciences 115, 10798 (2018).

[43] Redcay, E. & Schilbach, L. Using second-person neuroscience to elucidate the mechanisms of social interaction. Nature Reviews Neuroscience 20, 495–505 (2019).

[44] Müller-Pinzler, L., Krach, S., Krämer, U. M. & Paulus, F. M. The Social Neuroscience of Interpersonal Emotions, 241–256 (Springer International Publishing, Cham, 2017).

[45] Szymanski, C. et al. Teams on the same wavelength perform better: Inter-brain phase synchronization constitutes a neural substrate for social facilitation. NeuroImage 152, 425–436 (2017).

[46] Rice, K., Moraczewski, D. & Redcay, E. Perceived live interaction modulates the developing social brain. Social cognitive and affective neuroscience 11, 1354–1362 (2016).

[47] Anumanchipalli, G. K., Chartier, J. & Chang, E. F. Speech synthesis from neural decoding of spoken sentences. Nature 568, 493–498 (2019).

[48] Miller, K. J. A library of human electrocorticographic data and analyses. Nature Human Behaviour 3, 1225–1235 (2019).

[49] Takaura, K., Tsuchiya, N. & Fujii, N. Frequencydependent spatiotemporal profiles of visual responses recorded with subdural ecog electrodes in awake monkeys: Differences between high- and low-frequency activity. NeuroImage 124, 557–572 (2016).

[50] Gunduz, A. et al. Neural correlates of visual-spatial attention in electrocorticographic signals in humans. Frontiers in human neuroscience 5, 89–89 (2011).

[51] Pistohl, T., Ball, T., Schulze-Bonhage, A., Aertsen, A. & Mehring, C. Prediction of arm movement trajectories from ecog-recordings in humans. Journal of Neuroscience Methods 167, 105–114 (2008).

[52] Ball, T., Kern, M., Mutschler, I., Aertsen, A. & Schulze-Bonhage, A. Signal quality of simultaneously recorded invasive and non-invasive eeg. NeuroImage 46, 708–716 (2009).

[53] Kanth, S. T. & Ray, S. Electrocorticogram (ecog) is highly informative in primate visual cortex. The Journal of Neuroscience 40, 2430 (2020).

[54] Schalk, G. & Leuthardt, E. C. Brain-computer interfaces using electrocorticographic signals. IEEE Reviews in Biomedical Engineering 4, 140–154 (2011).

[55] Jacobs, J. & Kahana, M. J. Direct brain recordings fuel advances in cognitive electrophysiology. Trends in Cognitive Sciences 14, 162–171 (2010).

[56] Talakoub, O. et al. Reconstruction of reaching movement trajectories using electrocorticographic signals in humans. PLOS ONE 12, e0182542– (2017).

[57] Talakoub, O. et al. Temporal alignment of electrocorticographic recordings for upper limb movement. Frontiers in neuroscience 8, 431–431 (2015).

[58] Pistohl, T., Schulze-Bonhage, A., Aertsen, A., Mehring, C. & Ball, T. Decoding natural grasp types from human ecog. NeuroImage 59, 248–260 (2012).

[59] Chung, J. W. et al. Beta-band oscillations in the supplementary motor cortex are modulated by levodopa and associated with functional activity in the basal ganglia. NeuroImage: Clinical 19, 559–571 (2018).

[60] Peterson, S. M. & Ferris, D. P. Differentiation in theta and beta electrocortical activity between visual and physical perturbations to walking and standing balance. eneuro 5, ENEURO.0207–18.2018 (2018).

[61] Tan, H. et al. Decoding gripping force based on local field potentials recorded from subthalamic nucleus in humans. eLife 5, e19089 (2016).

[62] Milekovic, T., Truccolo, W., Grün, S., Riehle, A. & Brochier, T. Local field potentials in primate motor cortex encode grasp kinetic parameters. NeuroImage 114, 338–355 (2015).

[63] Ofori, E., Coombes, S. A. & Vaillancourt, D. E. 3d cortical electrophysiology of ballistic upper limb movement in humans. NeuroImage 115, 30–41 (2015).

[64] Gabriel, P. G. et al. Neural correlates of unstructured motor behaviors. Journal of Neural Engineering 16, 066026 (2019).

[65] Alasfour, A. et al. Coarse behavioral context decoding. Journal of Neural Engineering 16, 016021 (2019).

[66] Wang, N. X., Farhadi, A., Rao, R. P. & Brunton, B. W. AJILE movement prediction: Multimodal deep learning for natural human neural recordings and video. In Thirty-Second AAAI Conference on Artificial Intelligence (2018).

[67] Wang, N. X. R., Olson, J. D., Ojemann, J. G., Rao, R. P. N. & Brunton, B. W. Unsupervised Decoding of Long-Term, Naturalistic Human Neural Recordings with Automated Video and Audio Annotations. Frontiers in Human Neuroscience 10 (2016).

[68] Vansteensel, M. J. et al. Task-free electrocorticography frequency mapping of the motor cortex. Clinical Neurophysiology 124, 1169–1174 (2013).

[69] Chao, Z., Nagasaka, Y. & Fujii, N. Long-term asynchronous decoding of arm motion using electrocorticographic signals in monkey. Frontiers in Neuroengineering 3, 3 (2010).

[70] Mathis, A. et al. DeepLabCut: Markerless pose estimation of user-defined body parts with deep learning. Tech. Rep., Nature Publishing Group (2018).

[71] Singh, S. H., Peterson, S. M., Rao, R. P. N. & Brunton, B. W. Towards naturalistic human neuroscience and neuroengineering: behavior mining in long-term video and neural recordings. (2020).

[72] Brunton, B. W. & Beyeler, M. Data-driven models in human neuroscience and neuroengineering. Current Opinion in Neurobiology 58, 21–29 (2019).

[73] Datta, S. R., Anderson, D. J., Branson, K., Perona, P. & Leifer, A. Computational neuroethology: A call to action. Neuron 104, 11–24 (2019).

[74] Berman, G. J. Measuring behavior across scales. BMC Biology 16, 23 (2018).

[75] Brown, A. X. & de Bivort, B. Ethology as a physical science. Nature Physics 14, 653–657 (2018).

[76] Anderson, D. J. & Perona, P. Toward a science of computational ethology. Neuron 84, 18–31 (2014).

[77] Mooshagian, E., Wang, C., Holmes, C. D. & Snyder, L. H. Single units in the posterior parietal cortex encode patterns of bimanual coordination. Cerebral Cortex 28, 1549–1567 (2018).

[78] Ebner, T. J., Hendrix, C. M. & Pasalar, S. Past, present, and emerging principles in the neural encoding of movement. Advances in experimental medicine and biology 629, 127–137 (2009).

[79] Heldman, D. A., Wang, W., Chan, S. S. & Moran, D. W. Local field potential spectral tuning in motor cortex during reaching. IEEE Transactions on Neural Systems and Rehabilitation Engineering 14, 180–183 (2006).

[80] Donchin, O. et al. Single-unit activity related to bimanual arm movements in the primary and supplementary motor cortices. Journal of Neurophysiology 88, 3498–3517 (2002).

[81] Derix, J., Iljina, O., Schulze-Bonhage, A., Aertsen, A. & Ball, T. doctor or darling? decoding the communication partner from ecog of the anterior temporal lobe during non-experimental, real-life social interaction. Frontiers in Human Neuroscience 6, 251 (2012).

[82] Dumas, G., Nadel, J., Soussignan, R., Martinerie, J. & Garnero, L. Inter-brain synchronization during social interaction. PloS one 5, e12166 (2010).

[83] Farshchian, A. et al. Adversarial domain adaptation for stable brain-machine interfaces. (2018).

[84] Yang, Y., Chang, E. F. & Shanechi, M. M. Dynamic tracking of non-stationarity in human ecog activity. In 2017 39th Annual International Conference of the IEEE Engineering in Medicine and Biology Society (EMBC), 1660–1663 (IEEE, 2017).

[85] Klosterman, S. L., Estepp, J. R., Monnin, J. W. & Christensen, J. C. Day-to-day variability in hybrid, passive brain-computer interfaces: Comparing two studies assessing cognitive workload. In 2016 38th Annual International Conference of the IEEE Engineering in Medicine and Biology Society (EMBC), 1584–1590 (IEEE, 2016).

[86] Bigdely-Shamlo, N., Mullen, T., Kreutz-Delgado, K. & Makeig, S. Measure projection analysis: a probabilistic approach to eeg source comparison and multi-subject inference. NeuroImage 72, 287–303 (2013).

[87] Skarpaas, T. L., Tcheng, T. K. & Morrell, M. J. Clinical and electrocorticographic response to antiepileptic drugs in patients treated with responsive stimulation. Epilepsy & Behavior 83, 192–200 (2018).

[88] Struck, A. F., Cole, A. J., Cash, S. S. & Westover, M. B. The number of seizures needed in the emu. Epilepsia 56, 1753–1759 (2015).

[89] Engel, A. K. & Fries, P. Beta-band oscillations— signalling the status quo? Current Opinion in Neurobiology 20, 156–165 (2010).

[90] Tam, W.-k., Wu, T., Zhao, Q., Keefer, E. & Yang, Z. Human motor decoding from neural signals: a review. BMC Biomedical Engineering 1, 22 (2019).

[91] Branco, M. P., de Boer, L. M., Ramsey, N. F. & Vansteensel, M. J. Encoding of kinetic and kinematic movement parameters in the sensorimotor cortex: A brain-computer interface perspective. European Journal of Neuroscience 50, 2755–2772 (2019).

[92] Branco, M. P. et al. High-frequency band temporal dynamics in response to a grasp force task. Journal of Neural Engineering 16, 056009 (2019).

[93] Manning, J. R., Jacobs, J., Fried, I. & Kahana, M. J. Broadband shifts in local field potential power spectra are correlated with single-neuron spiking in humans. The Journal of Neuroscience 29, 13613 (2009).

[94] Başar, E., Başar-Eroglu, C., Karakaş, S. & Schürmann, M. Gamma, alpha, delta, and theta oscillations govern cognitive processes. International Journal of Psychophysiology 39, 241–248 (2001).

[95] van Kerkoerle, T. et al. Alpha and gamma oscillations characterize feedback and feedforward processing in monkey visual cortex. Proceedings of the National Academy of Sciences 111, 14332 (2014).

[96] Miller, K. J., Zanos, S., Fetz, E. E., den Nijs, M. & Ojemann, J. G. Decoupling the cortical power spectrum reveals real-time representation of individual finger movements in humans. The Journal of Neuroscience 29, 31–32 (2009).

[97] Lisberger, S. G. & Medina, J. F. How and why neural and motor variation are related. Current opinion in neurobiology 33, 110–116 (2015).

[98] Gomez-Marin, A., Paton, J. J., Kampff, A. R., Costa, R. M. & Mainen, Z. F. Big behavioral data: psychology, ethology and the foundations of neuroscience. Nature Neuroscience 17, 1455–1462 (2014).

[99] Melnik, A. et al. Systems, subjects, sessions: To what extent do these factors influence eeg data? Frontiers in human neuroscience 11, 150–150 (2017).

[100] Gliske, S. V. et al. Variability in the location of high frequency oscillations during prolonged intracranial eeg recordings. Nature Communications 9, 2155 (2018).

[101] Nurse, E. S. et al. Consistency of long-term subdural electrocorticography in humans. IEEE Transactions on Biomedical Engineering 65, 344–352 (2018).

[102] Gallego, J. A., Perich, M. G., Chowdhury, R. H., Solla, S. A. & Miller, L. E. Long-term stability of cortical population dynamics underlying consistent behavior. Nature Neuroscience 23 (2020).

[103] Liang, N. & Bougrain, L. Decoding finger flexion from band-specific ecog signals in humans. Frontiers in neuroscience 6, 91 (2012).

[104] Musall, S., Kaufman, M. T., Juavinett, A. L., Gluf, S. & Churchland, A. K. Single-trial neural dynamics are dominated by richly varied movements. Nature Neuroscience 22, 1677–1686 (2019).

[105] Ting, L. H. & McKay, J. L. Neuromechanics of muscle synergies for posture and movement. Current Opinion in Neurobiology 17, 622–628 (2007).

[106] Hu, K. et al. Decoding unconstrained arm movements in primates using high-density electrocorticography signals for brain-machine interface use. Scientific Reports 8, 10583 (2018).

[107] Georgopoulos, A., Schwartz, A. & Kettner, R. Neuronal population coding of movement direction. Science 233, 1416–1419. (1986).

[108] Gramfort, A. et al. Meg and eeg data analysis with mnepython. Frontiers in Neuroscience 7, 267 (2013).

[109] Gibbs, J. W. Fourier’s series. Nature 59, 606–606 (1899).

[110] Stolk, A. et al. Integrated analysis of anatomical and electrophysiological human intracranial data. Nature Protocols 13, 1699–1723 (2018).

[111] Oostenveld, R., Fries, P., Maris, E. & Schoffelen, J.-M. Fieldtrip: Open source software for advanced analysis of meg, eeg, and invasive electrophysiological data. Computational intelligence and neuroscience 2011, 156869–156869 (2011).

[112] Debnath, L. & Shah, F. A. Wavelet Transforms and Their Applications (Birkhäuser, 2015).

[113] Tzourio-Mazoyer, N. et al. Automated anatomical labeling of activations in spm using a macroscopic anatomical parcellation of the mni mri single-subject brain. NeuroImage 15, 273–289 (2002).

[114] Delorme, A. & Makeig, S. Eeglab: an open source toolbox for analysis of single-trial eeg dynamics including independent component analysis. Journal of Neuroscience Methods 134, 9–21 (2004).

[115] Peterson, S. M., Rios, E. & Ferris, D. P. Transient visual perturbations boost short-term balance learning in virtual reality by modulating electrocortical activity. Journal of neurophysiology 120 4, 1998–2010 (2018).

[116] Benjamini, Y. & Hochberg, Y. Controlling the false discovery rate: A practical and powerful approach to multiple testing. Journal of the Royal Statistical Society. Series B (Methodological) 57, 289–300 (1995).

[117] Master, S., Biase, N. d., Pedrosa, V. & Chiari, B. M. The long-term average spectrum in research and in the clinical practice of speech therapists. Pró-Fono Revista de Atualização Científica 18, 111–120 (2006).

[118] Huber, P. J. Robust estimation of a location parameter. The Annals of Mathematical Statistics 35, 73–101 (1964).

