## Supplementary Information for "Behavioral and Neural Variability of Naturalistic Arm Movements"

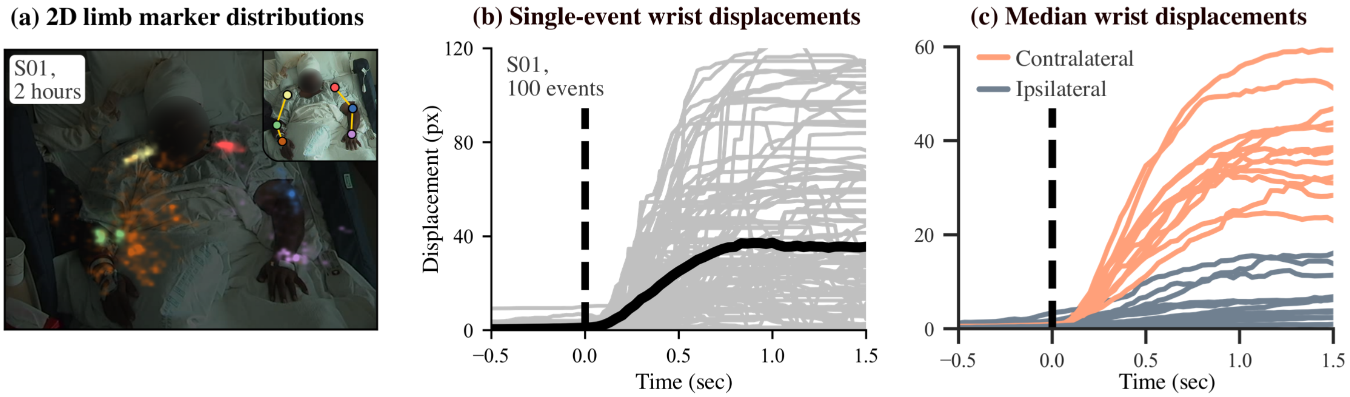


**Supplementary Fig. 1: Behavioral variability during naturalistic movements is large.** **(a)** An example of tracked joint markers during 2 hours of video monitoring for S01. Movement events of S01's right wrist are visualized, as the subject's electrodes were implanted in the left hemisphere. A heatmap of locations of each joint is shown. **(b)** An example of 100 randomly selected movement trajectories of the right wrist for S01, shown as displacement in pixels from the rest position, demonstrates the large variability seen across naturalistic arm movements. Solid black line denotes median displacement across events. Events are aligned by time of movement initiation (vertical dashed line). **(c)** Because we selected movement initiation events of the wrist contralateral to the hemisphere with implanted electrodes, the median contralateral wrist displacements (orange lines) across all 12 subjects are substantially greater than ipsilateral wrist displacements (gray lines).

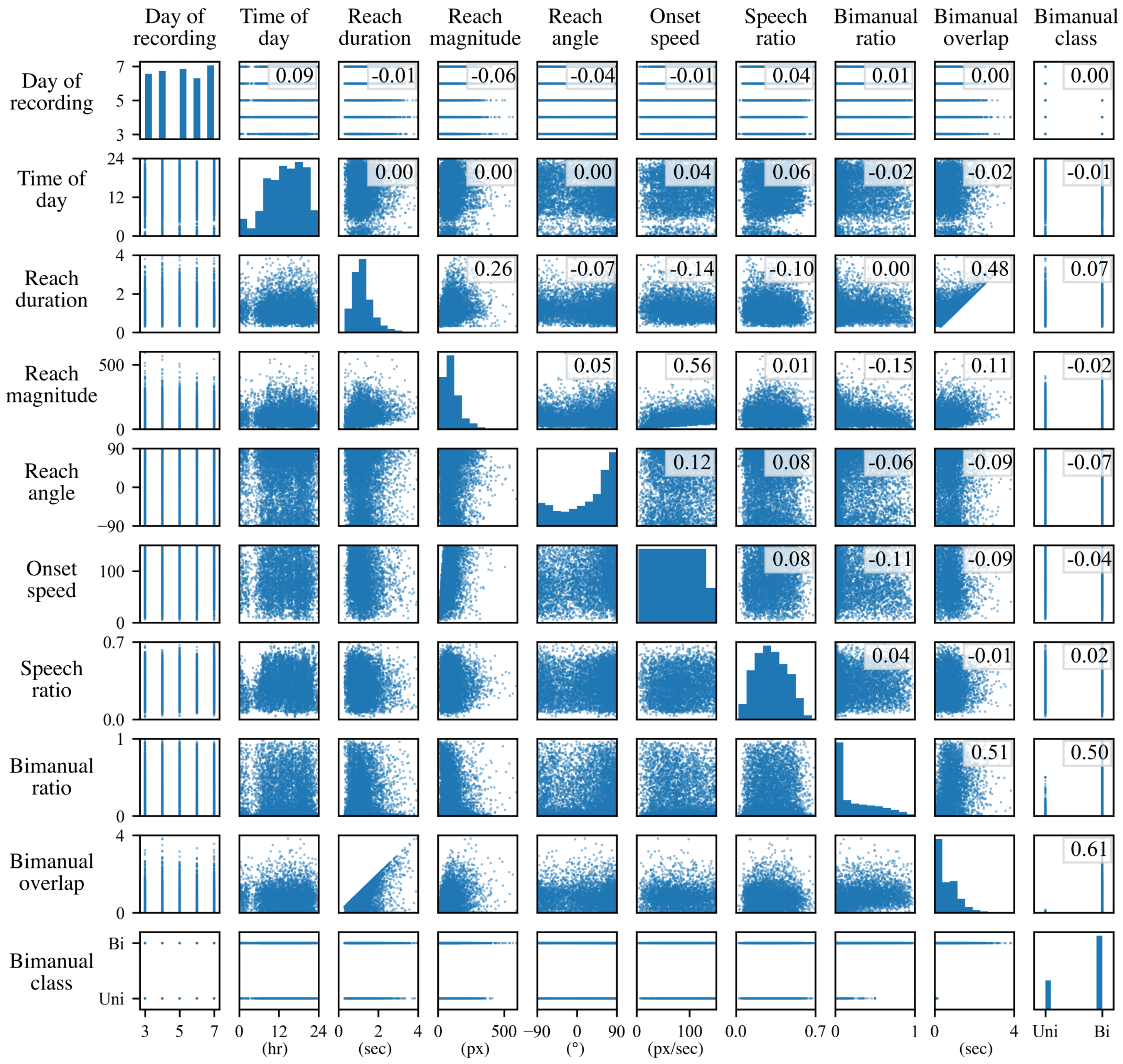


**Supplementary Fig. 2: Behavioral feature scatter matrix.** Group-level feature covariance is assessed using a scatter matrix, with feature pair Pearson correlation coefficients overlaid. Reach magnitude is positively correlated with reach duration and onset speed. Reach duration is positively correlated with bimanual overlap because the maximum amount of possible overlap depends on reach duration. All 3 bimanual features are highly correlated to each other as well.

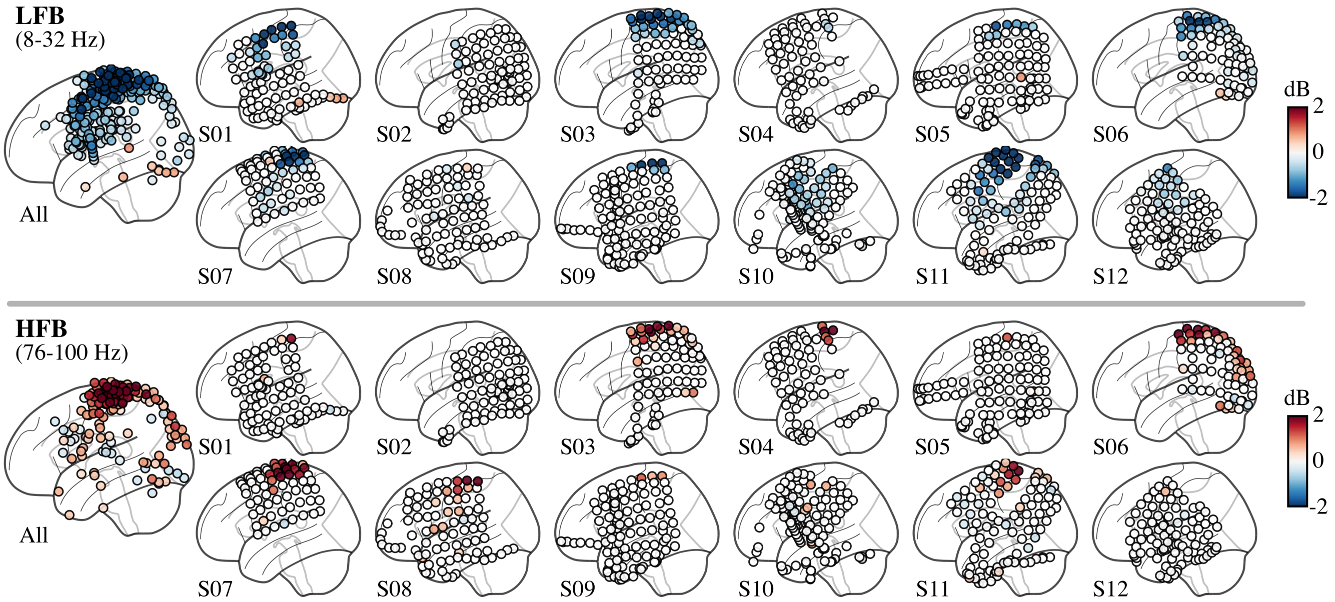


**Supplementary Fig. 3: Spectral power across electrodes for each subject.** Spectral power is shown averaged from 0 to 0.5 seconds across low/high frequency bands (LFB/HFB). Colors reflect median values across events. Median values that were not significantly different from -1.5 to -1 second baseline were set to zero (p>0.05 with false discovery rate correction, bootstrap statistics using 2000 permutations). The far left column shows spectral power for all 12 subjects; only the electrodes with significant power are shown.

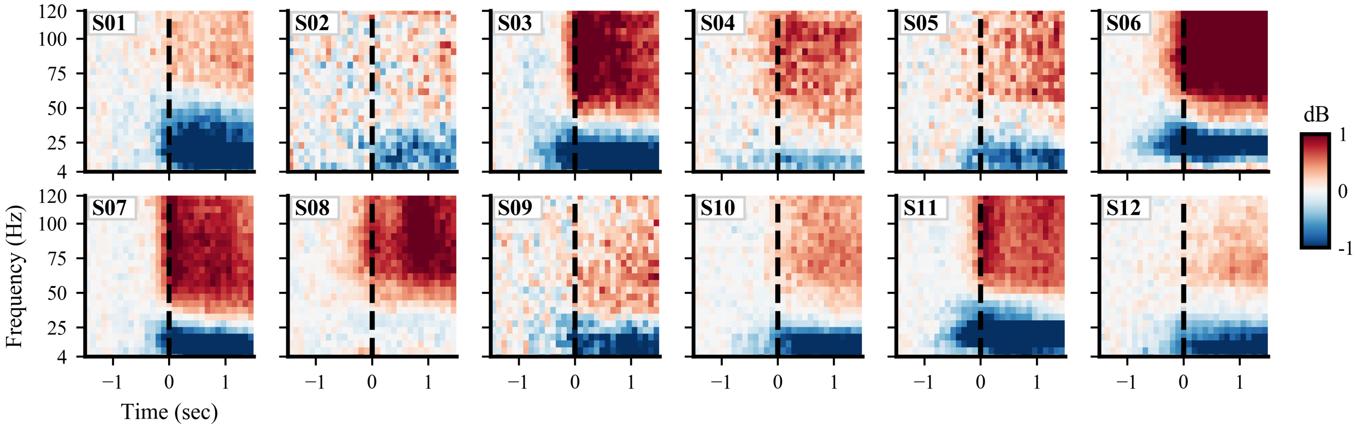


**Supplementary Fig. 4: Postcentral spectral power for each subject.** Postcentral median spectral power across contralateral arm movement onset events is shown for each subject. Colors indicate difference in spectral power relative to baseline (-1.5 to -1 sec) in dB. Movement onset is at time 0, denoted by dashed vertical line. No statistical masking is used.

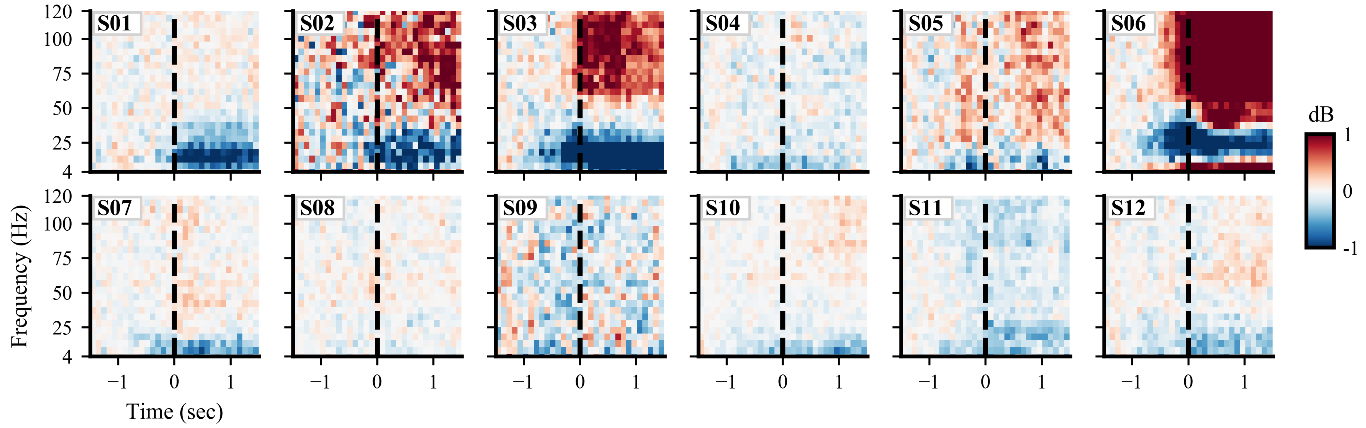


**Supplementary Fig. 5: Middle frontal spectral power for each subject.** Middle frontal median spectral power across contralateral arm movement onset events is shown for each subject. Colors indicate difference in spectral power relative to baseline (-1.5 to -1 sec) in dB. Movement onset is at time 0, denoted by dashed vertical line. No statistical masking is used.

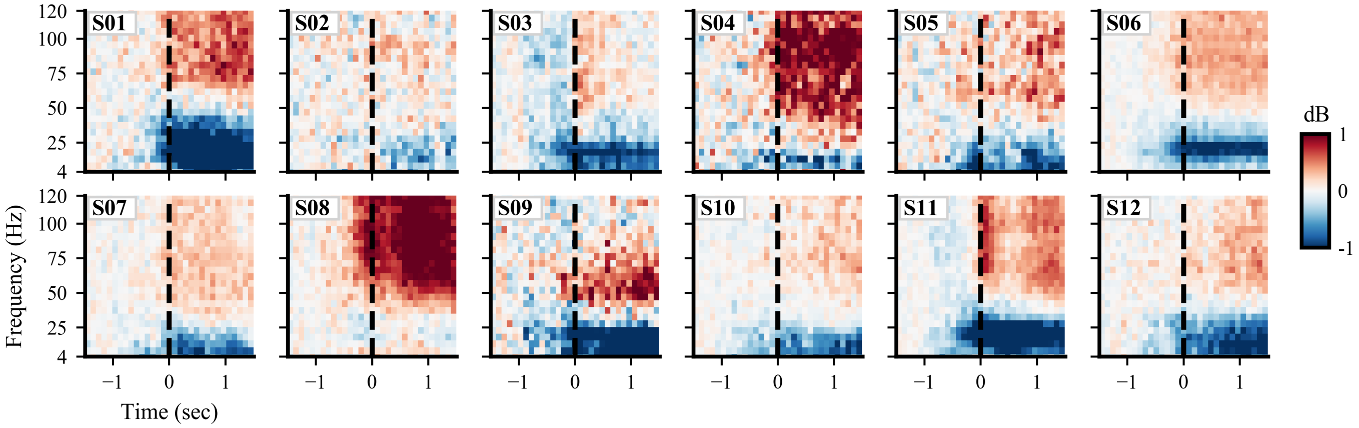


**Supplementary Fig. 6: Inferior parietal spectral power for each subject.** Inferior parietal median spectral power across contralateral arm movement onset events is shown for each subject. Colors indicate difference in spectral power relative to baseline (-1.5 to -1 sec) in dB. Movement onset is at time 0, denoted by dashed vertical line. No statistical masking is used.

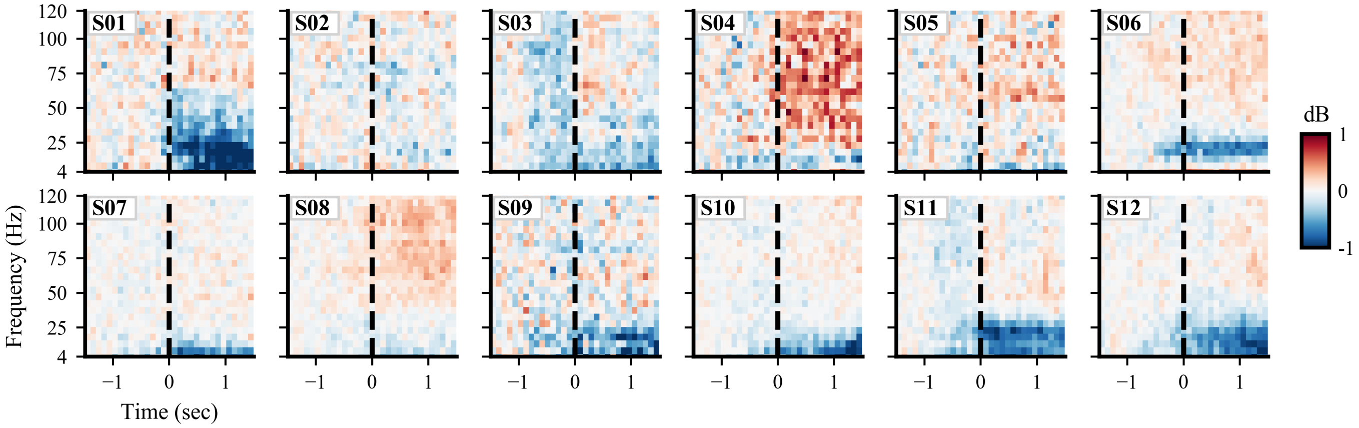


**Supplementary Fig. 7: Supramarginal spectral power for each subject.** Supramarginal median spectral power across contralateral arm movement onset events is shown for each subject. Colors indicate difference in spectral power relative to baseline (-1.5 to -1 sec) in dB. Movement onset is at time 0, denoted by dashed vertical line. No statistical masking is used.

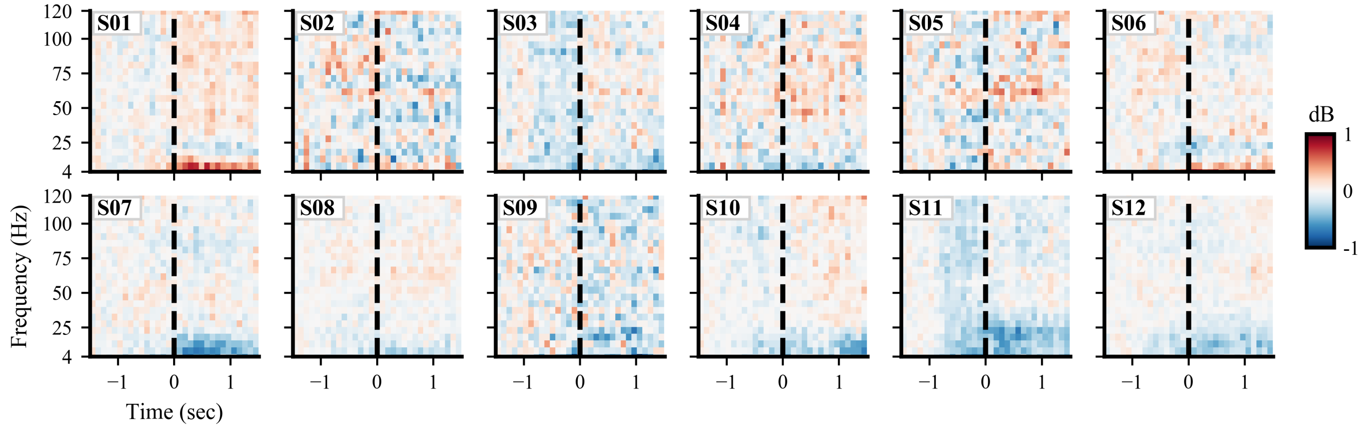


**Supplementary Fig. 8: Superior temporal spectral power for each subject.** Superior temporal median spectral power across contralateral arm movement onset events is shown for each subject. Colors indicate difference in spectral power relative to baseline (-1.5 to -1 sec) in dB. Movement onset is at time 0, denoted by dashed vertical line. No statistical masking is used.

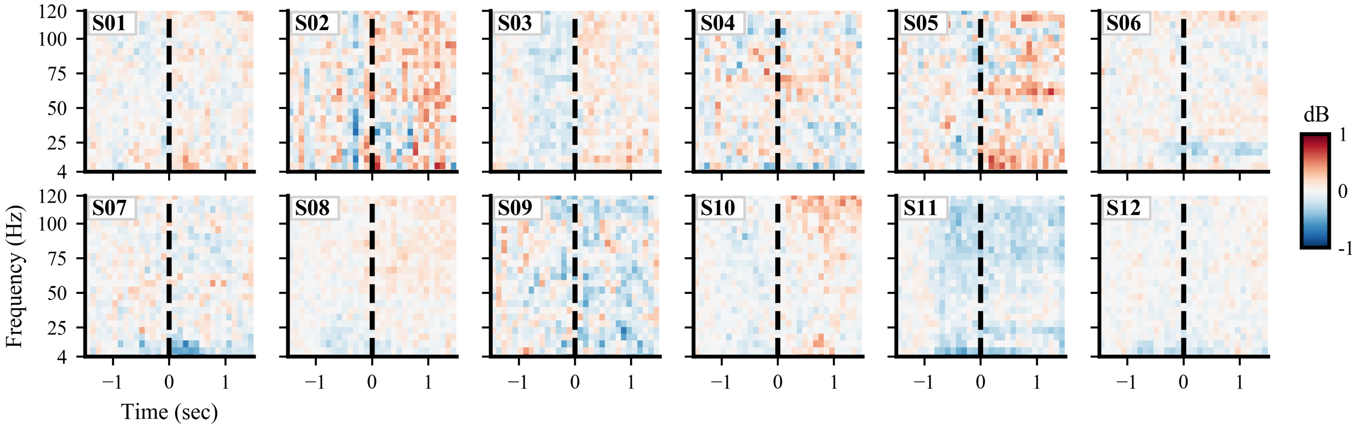


**Supplementary Fig. 9: Middle temporal spectral power for each subject.** Middle temporal median spectral power across contralateral arm movement onset events is shown for each subject. Colors indicate difference in spectral power relative to baseline (-1.5 to -1 sec) in dB. Movement onset is at time 0, denoted by dashed vertical line. No statistical masking is used.

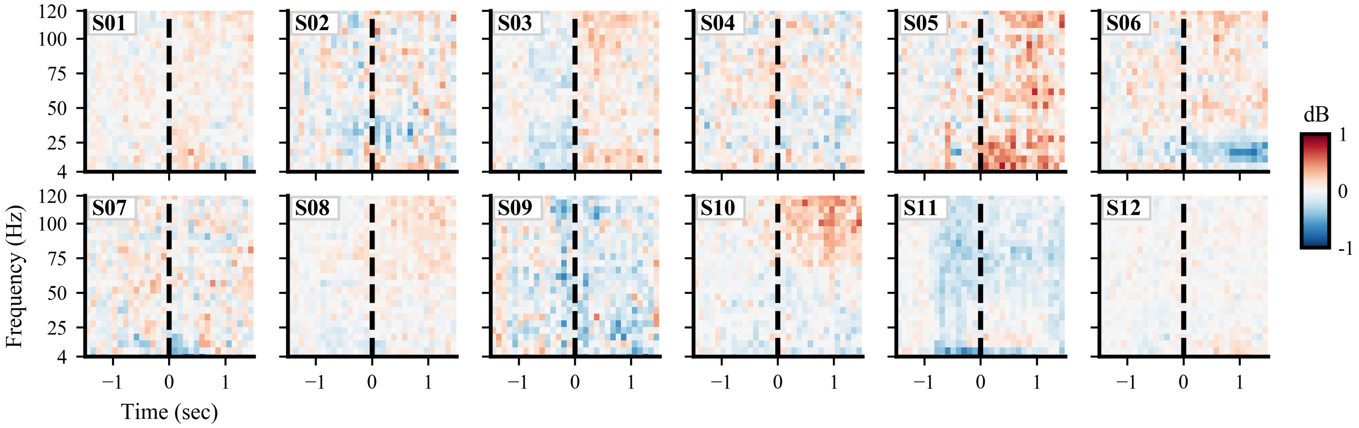


**Supplementary Fig. 10: Inferior temporal spectral power for each subject.** Inferior temporal median spectral power across contralateral arm movement onset events is shown for each subject. Colors indicate difference in spectral power relative to baseline (-1.5 to -1 sec) in dB. Movement onset is at time 0, denoted by dashed vertical line. No statistical masking is used.

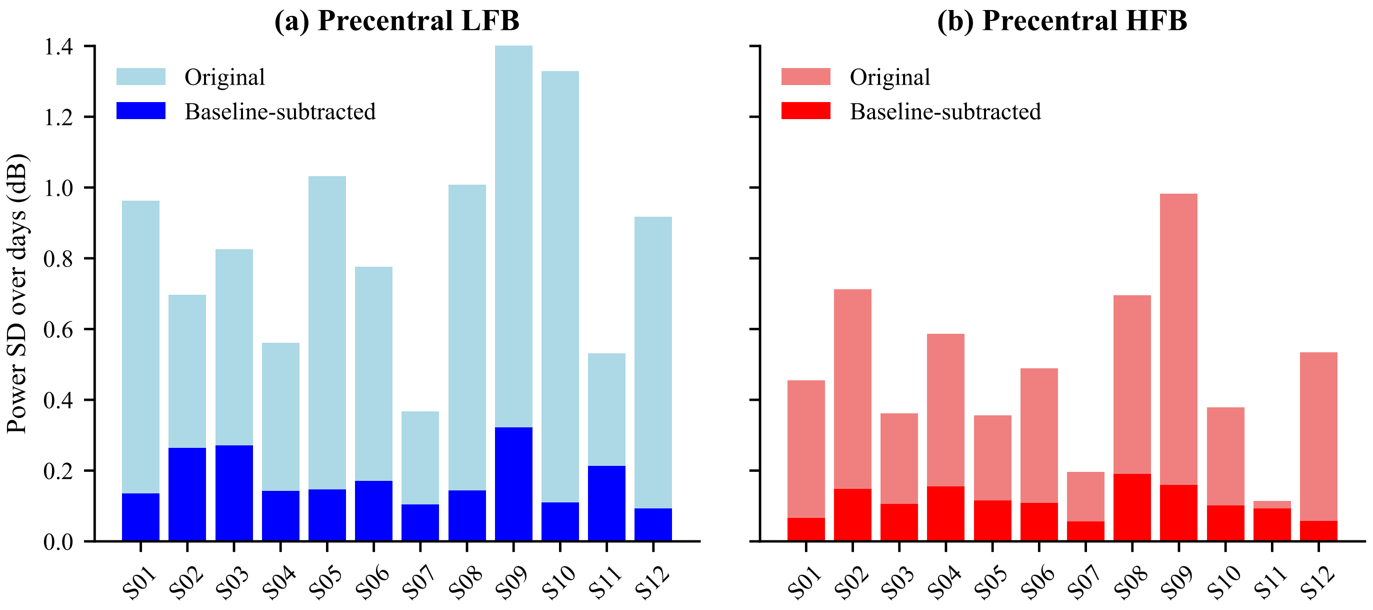


**Supplementary Fig. 11: Baseline subtraction minimizes the spectral power variability across recording days.** The standard deviation (SD) of average low frequency (8–32 Hz) and high frequency (76-100 Hz) band spectral power across recording days is shown with and without subtraction of a -1.5 to -1 second baseline. Baseline subtraction reduces the variation across recording days, as expected, but clear variability remains.

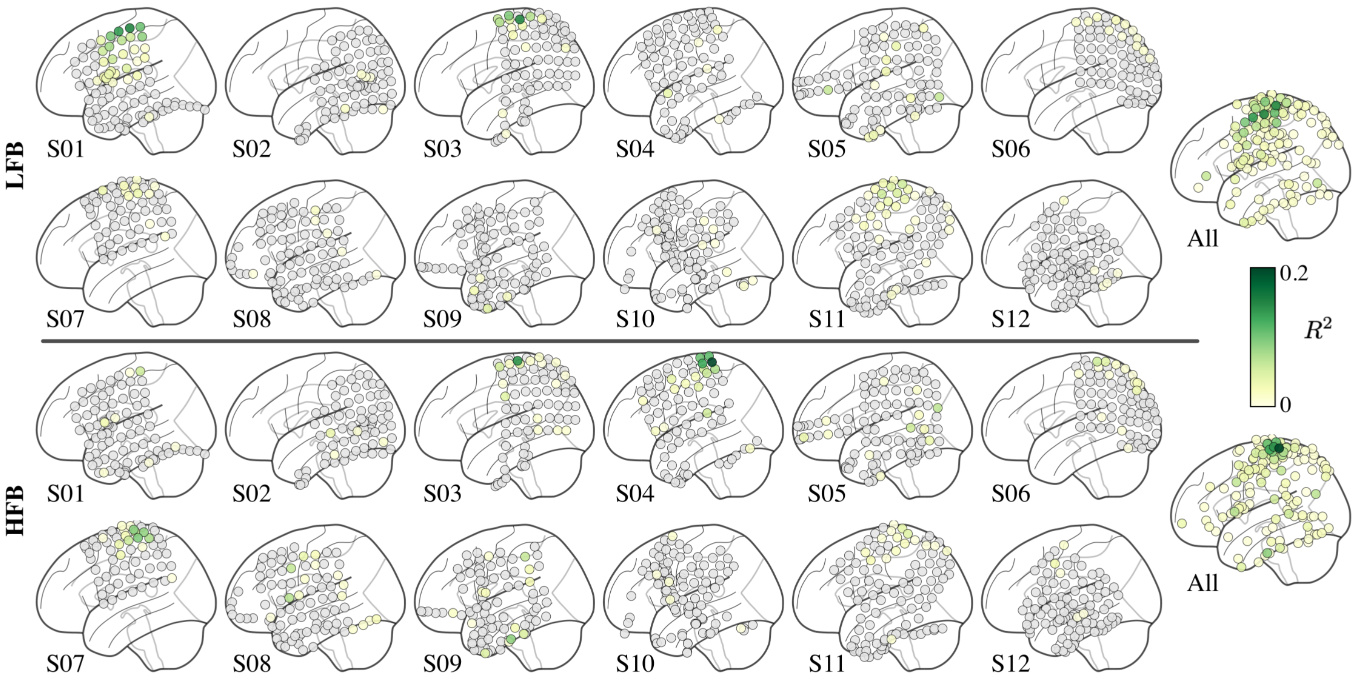


**Supplementary Fig. 12: Single-subject regression model fits.** Full model $R^2$ scores on withheld data are shown across subjects for low/high frequency band (LFB/HFB) spectral power features. The far right column displays electrodes across all subjects with $R^2>0$ averaged across 200 train/test splits. Well-fit models with the largest positive $R^2$ values are consistently located in fronto-parietal sensorimotor areas for most subjects.

| | | **Reach Duration** | **Reach Magnitude** | **Reach Angle** | **Onset Velocity** | **Speech Ratio** | **Bimanual Ratio** | **Bimanual Overlap** | **Bimanual Class** | **Day of Recording** | **Time of Day** |
|---|---|---|---|---|---|---|---|---|---|---|---|
| Precentral | LFB | 0.36 (0.25) | 0.24 (0.14) | 0.55 (0.30) | 0.30 (0.14) | 0.30 (0.19) | 0.41 (0.25) | 0.30 (0.21) | 0.30 (0.12) | 0.32 (0.07) | 0.25 (0.10) |
| | HFB | 0.38 (0.21) | 0.44 (0.22) | 0.55 (0.20) | 0.29 (0.15) | 0.36 (0.15) | 0.38 (0.18) | 0.22 (0.15) | 0.35 (0.13) | 0.28 (0.05) | 0.24 (0.07) |
| Middle Frontal | LFB | 0.41 (0.27) | 0.21 (0.24) | 0.46 (0.37) | 0.36 (0.20) | 0.28 (0.27) | 0.34 (0.25) | 0.20 (0.11) | 0.28 (0.15) | 0.26 (0.11) | 0.25 (0.11) |
| | HFB | 0.32 (0.24) | 0.41 (0.32) | 0.48 (0.22) | 0.26 (0.21) | 0.40 (0.23) | 0.36 (0.23) | 0.20 (0.23) | 0.37 (0.19) | 0.27 (0.09) | 0.26 (0.07) |
| Postcentral | LFB | 0.34 (0.17) | 0.32 (0.13) | 0.56 (0.26) | 0.24 (0.12) | 0.27 (0.14) | 0.43 (0.24) | 0.29 (0.16) | 0.28 (0.18) | 0.29 (0.06) | 0.22 (0.07) |
| | HFB | 0.39 (0.17) | 0.39 (0.13) | 0.53 (0.14) | 0.34 (0.09) | 0.33 (0.12) | 0.35 (0.09) | 0.30 (0.13) | 0.32 (0.12) | 0.28 (0.05) | 0.24 (0.04) |
| Inferior Parietal | LFB | 0.33 (0.25) | 0.33 (0.19) | 0.46 (0.27) | 0.28 (0.22) | 0.31 (0.12) | 0.42 (0.20) | 0.28 (0.25) | 0.19 (0.13) | 0.24 (0.09) | 0.21 (0.07) |
| | HFB | 0.25 (0.16) | 0.31 (0.22) | 0.38 (0.22) | 0.36 (0.15) | 0.32 (0.17) | 0.38 (0.12) | 0.22 (0.13) | 0.30 (0.12) | 0.27 (0.08) | 0.23 (0.09) |
| Supramarginal | LFB | 0.35 (0.21) | 0.33 (0.17) | 0.38 (0.25) | 0.30 (0.20) | 0.29 (0.19) | 0.39 (0.20) | 0.32 (0.18) | 0.22 (0.17) | 0.23 (0.09) | 0.22 (0.08) |
| | HFB | 0.33 (0.18) | 0.28 (0.13) | 0.38 (0.23) | 0.34 (0.20) | 0.25 (0.11) | 0.32 (0.22) | 0.29 (0.18) | 0.25 (0.09) | 0.25 (0.06) | 0.21 (0.05) |
| Superior Temporal | LFB | 0.37 (0.18) | 0.34 (0.16) | 0.34 (0.19) | 0.25 (0.14) | 0.26 (0.17) | 0.36 (0.21) | 0.35 (0.16) | 0.25 (0.17) | 0.26 (0.04) | 0.21 (0.06) |
| | HFB | 0.36 (0.19) | 0.25 (0.12) | 0.40 (0.20) | 0.29 (0.15) | 0.28 (0.12) | 0.20 (0.10) | 0.28 (0.15) | 0.27 (0.14) | 0.23 (0.06) | 0.23 (0.08) |
| Middle Temporal | LFB | 0.36 (0.13) | 0.28 (0.15) | 0.34 (0.23) | 0.22 (0.12) | 0.31 (0.18) | 0.31 (0.19) | 0.33 (0.13) | 0.28 (0.15) | 0.26 (0.06) | 0.20 (0.05) |
| | HFB | 0.25 (0.14) | 0.27 (0.12) | 0.33 (0.17) | 0.28 (0.14) | 0.29 (0.13) | 0.24 (0.08) | 0.25 (0.13) | 0.27 (0.17) | 0.24 (0.07) | 0.22 (0.07) |
| Inferior Temporal | LFB | 0.32 (0.11) | 0.29 (0.18) | 0.33 (0.19) | 0.22 (0.16) | 0.29 (0.21) | 0.31 (0.14) | 0.29 (0.16) | 0.28 (0.17) | 0.26 (0.06) | 0.19 (0.07) |
| | HFB | 0.31 (0.14) | 0.30 (0.12) | 0.26 (0.12) | 0.28 (0.17) | 0.26 (0.11) | 0.29 (0.13) | 0.29 (0.15) | 0.32 (0.17) | 0.24 (0.05) | 0.24 (0.10) |

**Supplementary Table 1: Reach angle is the most consistently retained feature for regression models in sensorimotor areas.** The average probability of retaining each feature across subjects following forward selection feature pruning is shown, projected to the 8 regions of interest (standard deviation is in parentheses). Regression models were fit to single-event low-frequency band (LFB) or high-frequency band (HFB) spectral power features. Reach angle is the feature most often retained, followed by reach duration, reach magnitude, bimanual ratio, and bimanual overlap.

| Subject | Recording days used | Hemisphere implanted | # surface electrodes | # depth electrodes |
|---|---|---|---|---|
| 01 | 4 | L | 86 | 8 |
| 02 | 4 | R | 70 | 16 |
| 03 | 4 | L | 80 | 0 |
| 04 | 5 | R | 84 | 0 |
| 05 | 3 | R | 106 | 0 |
| 06 | 5 | L | 80 | 0 |
| 07 | 5 | R | 64 | 0 |
| 08 | 5 | R | 92 | 0 |
| 09 | 5 | L | 98 | 28 |
| 10 | 5 | L | 86 | 40 |
| 11 | 5 | L | 106 | 0 |
| 12 | 5 | L | 92 | 24 |

**Supplementary Table 2: Single-subject recording days used and electrode information.** Surface electrodes refer to grid and strip electrodes placed on the cortical surface, while depth electrodes reach deep cortical and subcortical areas.
